# Evidence for cryptic gene flow in parthenogenetic stick insects of the genus *Timema*

**DOI:** 10.1101/2023.01.21.525009

**Authors:** Susana Freitas, Darren J. Parker, Marjorie Labédan, Zoé Dumas, Tanja Schwander

## Abstract

Obligately parthenogenetic species are expected to be short lived since the lack of sex and recombination should translate into a slower adaptation rate and increased accumulation of deleterious alleles. Some, however, are thought to have been reproducing without males for millions of years. It is not clear how these old parthenogens can escape the predicted long-term costs of parthenogenesis, but an obvious explanation is cryptic sex.

In this study we screen for signatures of cryptic sex in eight populations of four parthenogenetic species of *Timema* stick insects, some estimated to be older than 1M yrs. Low genotype diversity, homozygosity of individuals and high linkage disequilibrium (LD) unaffected by marker distances support exclusively parthenogenetic reproduction in six populations. However, in two populations (namely, of the species *Timema douglasi* and *T. monikensis*) we find strong evidence for cryptic sex, most likely mediated by rare males. These populations had comparatively high genotype diversities, lower LD, and a clear LD decay with genetic distance. Rare sex in species that are otherwise largely parthenogenetic could help explain the unusual success of parthenogenesis in the *Timema* genus and raises the question whether episodes of rare sex are in fact the simplest explanation for the persistence of many old parthenogens in nature.

## INTRODUCTION

Although sex is the most prevalent mode of reproduction in animals, many species are able to reproduce via female-producing parthenogenesis. These species can reproduce without males, either exclusively (obligate parthenogens), or facultatively (facultative and cyclical parthenogens) (Bell 1982). Parthenogenesis is associated with considerable short-term advantages relative to sexual reproduction, the two main ones being reproductive assurance (since parthenogenetic organisms only require one individual for reproduction) (Gerritsen 1980) and demographic advantage (since parthenogenic reproduction produces only females) (Williams 1975; Maynard Smith 1978). However, these short-term advantages are predicted to be coupled with long-term costs, including the accumulation of deleterious alleles (Muller 1964; Gabriel et al. 1993), and a slower rate of adaptive evolution (Hill and Robertson 1966; Barton 1995). These long-term costs are believed to result in the eventual extinction of obligately parthenogenetic species (Maynard Smith 1978; Gabriel et al. 1993).

Despite the expected long-term costs of parthenogenesis, several parthenogenetic species are thought to have been reproducing solely parthenogenetically for millions of years (Judson and Normark 1996; Schurko et al. 2009). This raises the question of how these old parthenogens have managed to escape the predicted long-term consequences of parthenogenesis thus far. One possible explanation would be that besides reproducing parthenogenetically, they occasionally reproduce sexually. Indeed, bouts of rare sex in mainly parthenogenetically reproducing populations would be sufficient to overcome many of the long-term costs of parthenogenesis and could prevent the mutational deterioration of otherwise clonal lineages (Green and Noakes 1995; Bengtsson 2009; D’Souza and Michiels 2010). Consistent with this idea, several species originally thought to be obligately parthenogenetic have been recently shown to exchange genetic material between individuals (cryptic gene flow), including the brine shrimp *Artemia parthenogenetica* (Boyer et al. 2021), and the bdelloid rotifers *Adineta vaga* (Vakhrusheva et al. 2020) and *Macrotrachella quadricornifera* (Signorovitch et al. 2015; Laine et al. 2022).

In bdelloid rotifers, males have never been observed (Birky 2010), however patterns of allele sharing between individuals are compatible with some form of rare or non-canonical sex (Signorovitch et al. 2015; Vakhrusheva et al. 2020; Laine et al. 2022). On the other hand, in *Artemia* brine shrimps it has been demonstrated that cryptic gene flow is mediated by rare males (Boyer et al. 2021). Similar to *Artemia*, rare males are documented in many obligately parthenogenetic species (Simon et al. 1991; Butlin et al. 1998; Delmotte et al. 2001; Tvedte et al. 2020), but they typically do not appear to reproduce sexually with parthenogenetic females (reviewed in (van der Kooi and Schwander 2014). In some species, males produced by parthenogenetic females can mate with females of related sexual strains, but while such matings can result in new parthenogenetic lineages via contagious parthenogenesis (Innes and Hebert 1988; Maccari et al. 2014), they do not mediate gene flow within parthenogenetic lineages. Nevertheless, formal tests of cryptic gene flow within parthenogenetic lineages remain scarce, and rare males may therefore mediate cryptic gene flow in a broader panel of “obligately” parthenogenetic species than generally assumed.

In this study, we assess whether cryptic gene flow occurs in parthenogenetic species of the stick insect genus *Timema* (Phasmatodea). This genus comprises five described parthenogenetic species (Sandoval and Vickery 1996; Vickery and Sandoval 1999, 2001) which represent at least seven independently-derived parthenogenetic lineages (Schwander et al. 2011). These lineages vary in age, and the oldest parthenogenetic *Timema* is thought to have been reproducing via parthenogenesis for over 1 million years (Schwander et al. 2011). Parthenogenetic *Timema* are very homozygous as they most likely reproduce via a form of automictic parthenogenesis where heterozygosity is completely lost each generation (Jaron et al. 2022; Larose et al. 2022). Rare males have been found in four of the five described parthenogenetic *Timema* species (Vickery and Sandoval 1999, 2001; Schwander et al. 2013), and these rare “parthenogenetic” males present normal male reproductive organs, and can mate and produce offspring with sexual sister-species females, although with reduced fertility (Schwander et al. 2013). It is not clear how such males are produced, however they are thought to originate from the accidental loss of an X chromosome (aneuploidy) during parthenogenesis (Schwander et al. 2013). Given the XX/X0 mechanism of sex determination in *Timema* (Schwander and Crespi 2009), if an XX parthenogenetic egg loses one X it could potentially develop into a male, a mechanism documented in other parthenogenetic insect species (Pijnacker and Ferwerda 1980; Blackman and Hales 1986).

To test for cryptic gene flow in *Timema* we use RADseq genotyping to estimate linkage disequilibrium (LD) in eight populations of parthenogenetic species. A population specific approach is appropriate for studies with these species, as they live in patchy habitats and have very limited dispersal. LD can serve as a measure of genetic exchange as with strict parthenogenesis alleles will be transmitted in a single block. Without sex or recombination LD will only be broken by mutation or allele conversion, and is thus expected to be high. Sexual events (with recombination or segregation), on the other hand, will decrease LD, even when they are very rare. While we find no evidence for cryptic gene flow in six populations, we show that in two of the eight studied populations of parthenogenetic *Timema*, LD within and between chromosomes is very low, and there is a consistent LD decay along the chromosomes. In one of these populations we also find several individuals with unusually high heterozygosity, indicative of recent sexual events. The results presented here are the first evidence for cryptic gene flow, most likely mediated by rare males, in “obligately” parthenogenetic *Timema* and contribute to the growing evidence that rare or non-canonical sex might have to be considered when studying the long-term persistence of (mostly) parthenogenetically reproducing species.

## MATERIAL AND METHODS

### Samples and Sequencing

We looked for evidence of gene flow in two populations from each of four parthenogenetic species of *Timema* (*T. genevievae, T. shepardi, T. monikensis, T. douglasi*), in a total of eight populations. A population of the sexual species *T. cristinae* was also included to establish a sexual reference for LD and heterozygosity patterns based on RADseq genotypes. We used a population specific approach because *Timema* are wingless and it has been estimated that they can travel a maximum of 128 metres per generation (Sandoval 2000), which is much less than the distance between the two studied populations in each parthenogenetic species (6 km, 31 km, 43 km and 219 km apart on a straight line for *T. monikensis, T. shepardi, T. douglasi* and *T. genevievae* respectively).

We genotyped 24 females for each of the eight parthenogenetic populations (192 females in total) and 24 individuals (12 females, 12 males) for the sexual population (see also Table S1). Additionally, because we found evidence for recent gene flow in one population of *T. monikensis* (see below) we included additional samples of *T. monikensis* from different collection years for both populations under study, which included four males from the FS population (Table S1).

DNA was extracted from the head or legs (Table S1) using the Qiagen DNeasy Blood and Tissue Kit, following manufacturer instructions. ddRAD libraries were prepared following the protocol from (Brelsford et al. 2016), with the enzymes EcoRI and MseI, and a 200-450 bp size selection after addition of Illumina TruSeq indexes. For logistic reasons, a few samples underwent a different size selection (300-500bp). 96 samples were pooled in each library thanks to barcoded adapters (see protocol in (Brelsford et al. 2016)). Libraries were multiplexed by pairs on two Illumina lanes using Illumina TruSeq indexes iA06 or iA12. 150bp Single-end sequencing was performed using Illumina Hiseq 2500 at the Lausanne Genomic Technologies Facility. Reads are available under BioProject accession number PRJNA798556.

### Reads processing

All scripts used in this pipeline are available online (https://github.com/SusanaNFreitas/cryptic_gene_flow). We used the STACKS software (version 2.5) (Catchen et al. 2013) to de-multiplex the reads with the “process radtags” module from the STACKS suite. Reads were trimmed from adaptors and filtered for minimum length (80 bp) with Cutadapt 2.3 (Martin 2011) and aligned against the corresponding reference genomes (Jaron et al. 2022) with BWA. Sam files were sorted and converted to bam using samtools 1.4 excluding multiple alignments and PHRED quality score of 30 or less. FreeBayes (version 1.3.3) (Garrison and Marth 2012) was used to call SNPs, which were then filtered by read depth (with GATK - version 4.2.0.0 (Rimmer et al. 2014), keeping only positions with DP > 8 and DP < 200), genotype quality (using a custom python script, keeping SNP genotypes with QUAL > 30), and missing data (with vcftools (Danecek et al. 2011), only positions with 75% data kept). The *vcfallelicprimitives* function of vcflib (Garrison et al. 2022) was used to break MNPs in the vcf file into SNPs, and only biallelic SNPs were kept (eliminating indels and multi-allelic positions with custom unix oneliners). Low coverage samples were excluded from further analyses (Table S1).

### Population Genetics

The total number of SNPs per species was estimated with the R package ADEGENET 2.1.3 (Jombart 2008; Jombart and Ahmed 2011). Genetic distances were estimated using the Euclidean distance with the dist() function. Linkage Disequilibrium (LD) was estimated with plink 1.9 (Purcell et al. 2007), the *r*^2^ value estimated within and between linkage groups with options --*r2* and --*inter-chr*, and LD decay within linkage group with options --*r2, --ld-window 999999* and *--ld-window-kb* 8000. Only SNPs with minor allele frequency > 0.2 were used to estimate *r*^2^. For LD decay, we only used SNPs with available chromosomal coordinates (Jaron et al. 2022; Parker et al. 2022), which were based on chromosomal information from the species *T. cristinae* (assembly v1.3 (Nosil et al. 2018)). The best fit line estimated on the LD decay plots was generated with the geom_smooth() function using the gam method, or the loess method whenever fewer than 1000 data points were available (chromosomal interactions). Because the X chromosome was misassembled in the v1.3*T. cristinae* assembly, we used the positional information for SNPs on all chromosomes except the X (Jaron et al. 2022; Parker et al. 2022). *r*^2^ values per population were plotted with the package ggplot2 (Wickham 2016) in R. Relative heterozygosity was estimated for each individual by dividing the number of heterozygous positions by the total number of called polymorphic positions.

## RESULTS

### Genetic diversity patterns – sexual vs parthenogenetic

Sexual *Timema* populations, as exemplified by the *T. cristinae* population included here, present high relative heterozygosity and genotype diversity (Figure S1, see also (Jaron et al. 2022). As expected for a sexual population, where recombination and segregation of chromosomes break LD, *r*^2^ values both within and between chromosomes were very low (Figure 1 - A), even between SNPs separated by small distances (Figure 1 - B).

**Figure 1.**
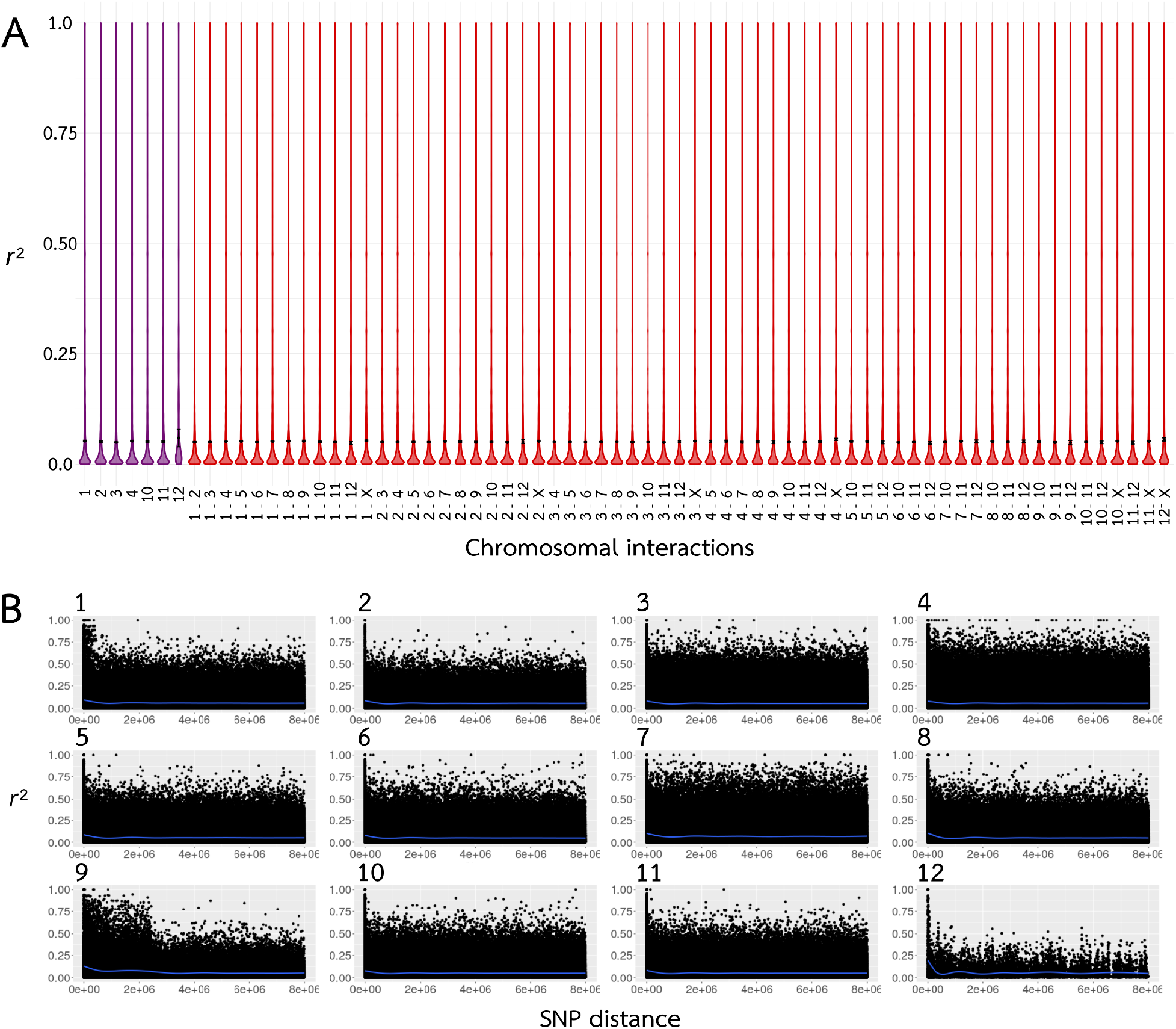
Linkage Disequilibrium (LD) in the sexual species *T. cristinae*. **A**. Violin plots of the Linkage Disequilibrium (LD) values within (purple) and between (red) chromosomes (linkage groups). Black dot represents the median LD (*r*^2^). Note LD values for chromosomes 5 to 9 were omitted from the plot for better readability (with all chromosomes presenting similar LD) **B**. LD decay for each linkage group. LD decay was estimated for all chromosomes excluding the X, for which we did not have positional information (see methods). A line of best fit is shown in blue.

In contrast to the sexual population with high genotype diversity, populations reproducing solely by parthenogenesis are prone to undergo recurrent sweeps. This should result in the fixation of one or few clones (genotypes), with minor genetic differences between individuals within clonal lineages. In *Timema*, strict parthenogenesis would further be associated with very low heterozygosity for all individuals in a population, since their parthenogenesis mechanism leads to the complete or largely complete loss of heterozygosity every generation (Jaron et al. 2022; Larose et al. 2022).

On the other hand, if parthenogenetic *Timema* populations are undergoing rare sex we would expect to see an increase in the genotypic diversity, and perhaps some individuals with elevated heterozygosity, which would indicate that they were produced via sex. From the RADseq reads mapped to each species’ genome, we recovered 25777, 17997, 14326 and 10617 SNPs for *T. genevievae, T. monikensis, T. douglasi* and *T. shepardi*, respectively. These were then used to infer the number of clones in our populations by estimating pairwise genetic distances between individuals, visualised as neighbour-joining networks (Figure 2). To explicitly look for cryptic sex, we then measured LD and LD decay in each of the eight parthenogenetic populations (Figures 3-5, and S2-S9) and estimated relative heterozygosity for each individual (Figure 6).

**Figure 2.**
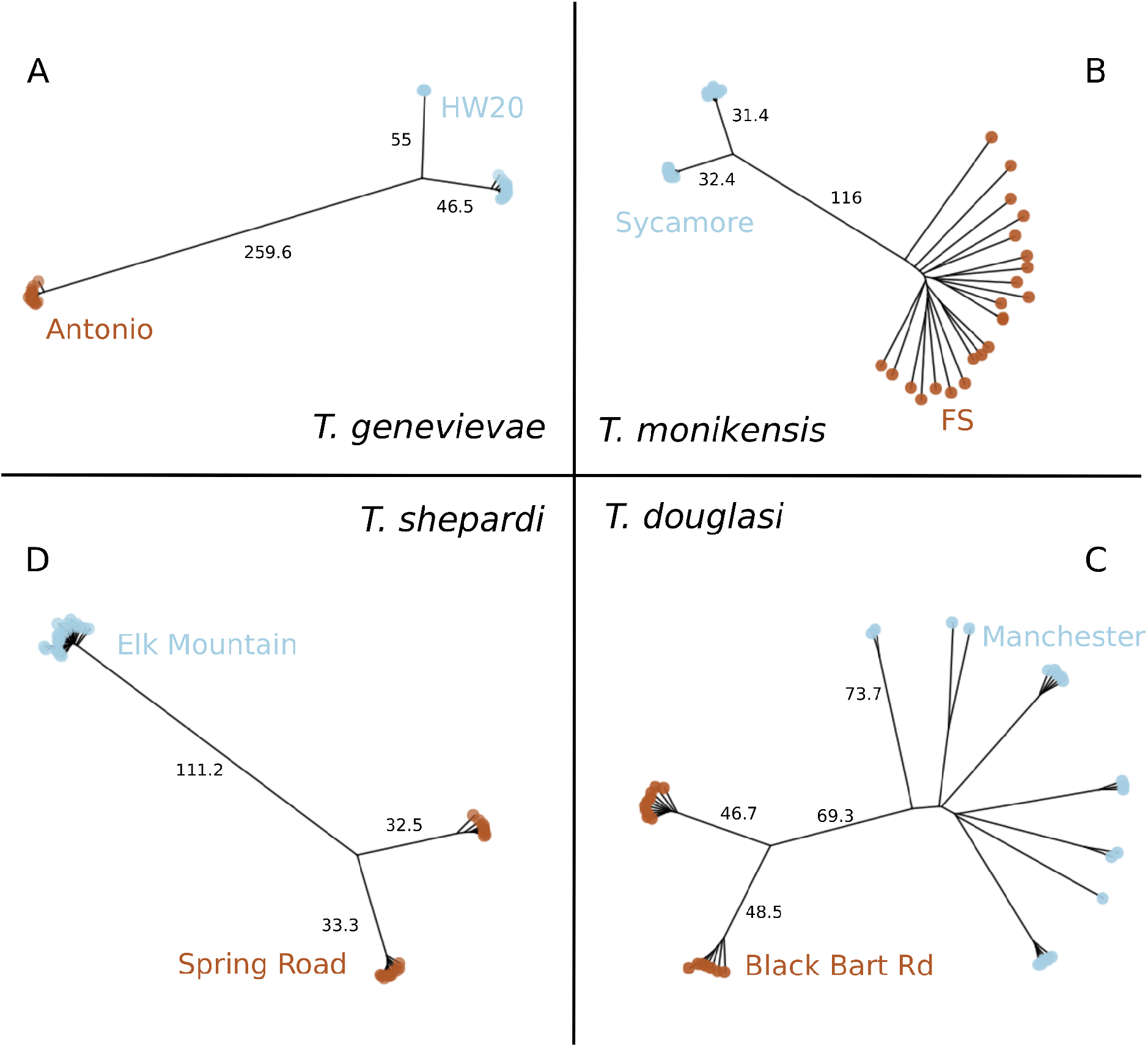
Neighbour-joining networks of pairwise distances between individuals within parthenogenetic *Timema* species: **A**. *T. genevievae*. **B**. *T. monikensis*. **C**. *T. douglasi*. **D**. *T. shepardi*. Individuals of the two studied populations in each species are distinguished by brown and blue labels.

We found two distinct patterns in the genetic diversity estimates (pairwise distances, LD and heterozygosity) among the populations of the parthenogenetic species. In six of the eight populations we found no evidence for cryptic sex, and the observed patterns clearly matched the expectations for a purely parthenogenetic population: genotype diversities were very low (one or two clones per population, Figure 2), and LD was high (Figures 3-A, S2-3, S4-A and S5-A), and did not decay over increasing genetic distances (Figures 3-B, S2-3, S6-7, S8-A and S9 - A). We found clear evidence for gene flow in the remaining two populations, with high diversity of genotypes (Figure 2), low LD (Figure 4) and evidence of LD decay (Figure 5).

**Figure 3.**
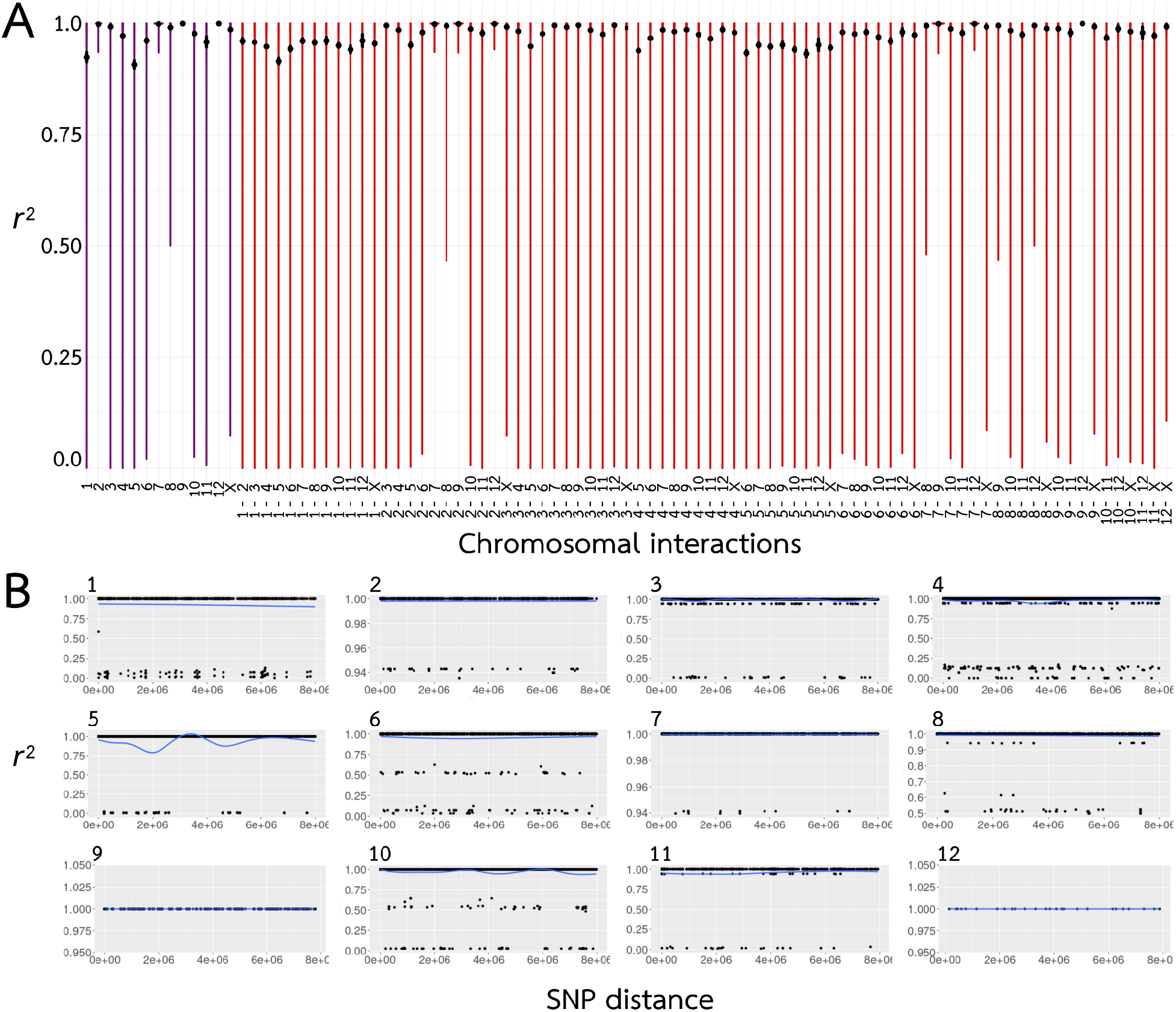
Linkage Disequilibrium (LD) in the parthenogenetic species *T. genevievae* (population HW20). **A**. Violin plots of the Linkage Disequilibrium (LD) values within (purple) and between (red) chromosomes (linkage groups). Black dot represents the median LD (*r*^2^). **B**. LD decay for each linkage group. LD decay was estimated for all chromosomes excluding the X, for which we did not have positional information (see methods). A line of best fit is shown in blue. LD values for *T. genevievae* - population HW20 are presented here as an illustrative example, but see Figures S2-S9 for all parthenogenetic populations.

**Figure 4.**
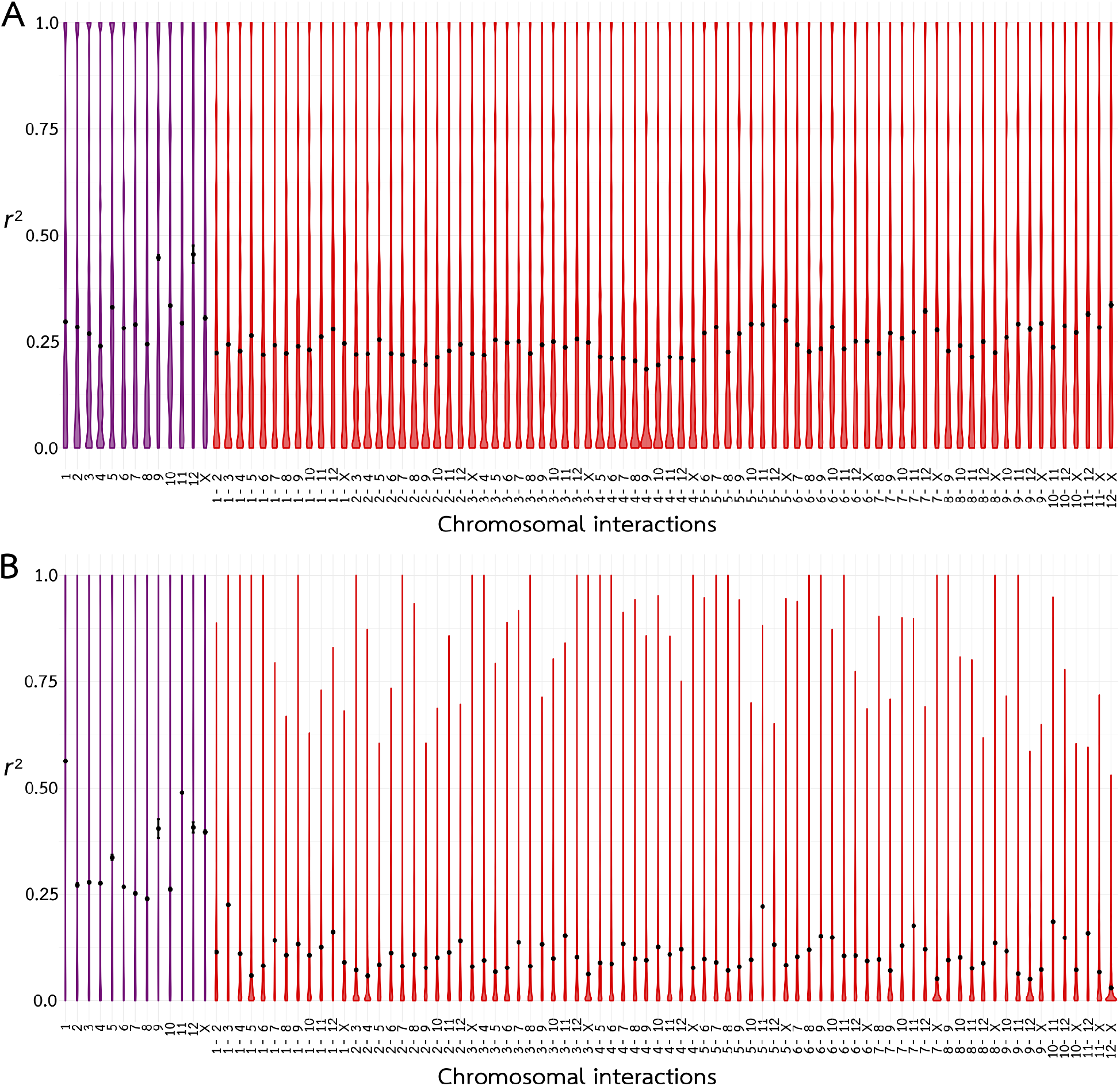
Linkage Disequilibrium (LD) values in parthenogenetic populations showing evidence for gene flow. LD values within (purple) and between (red) chromosomes (linkage groups) represented in violin plots, with the black dot representing the median LD (*r*^2^). **A**. *T. douglasi*, population Manchester. **B**. *T. monikensis*, population FS.

**Figure 5.**
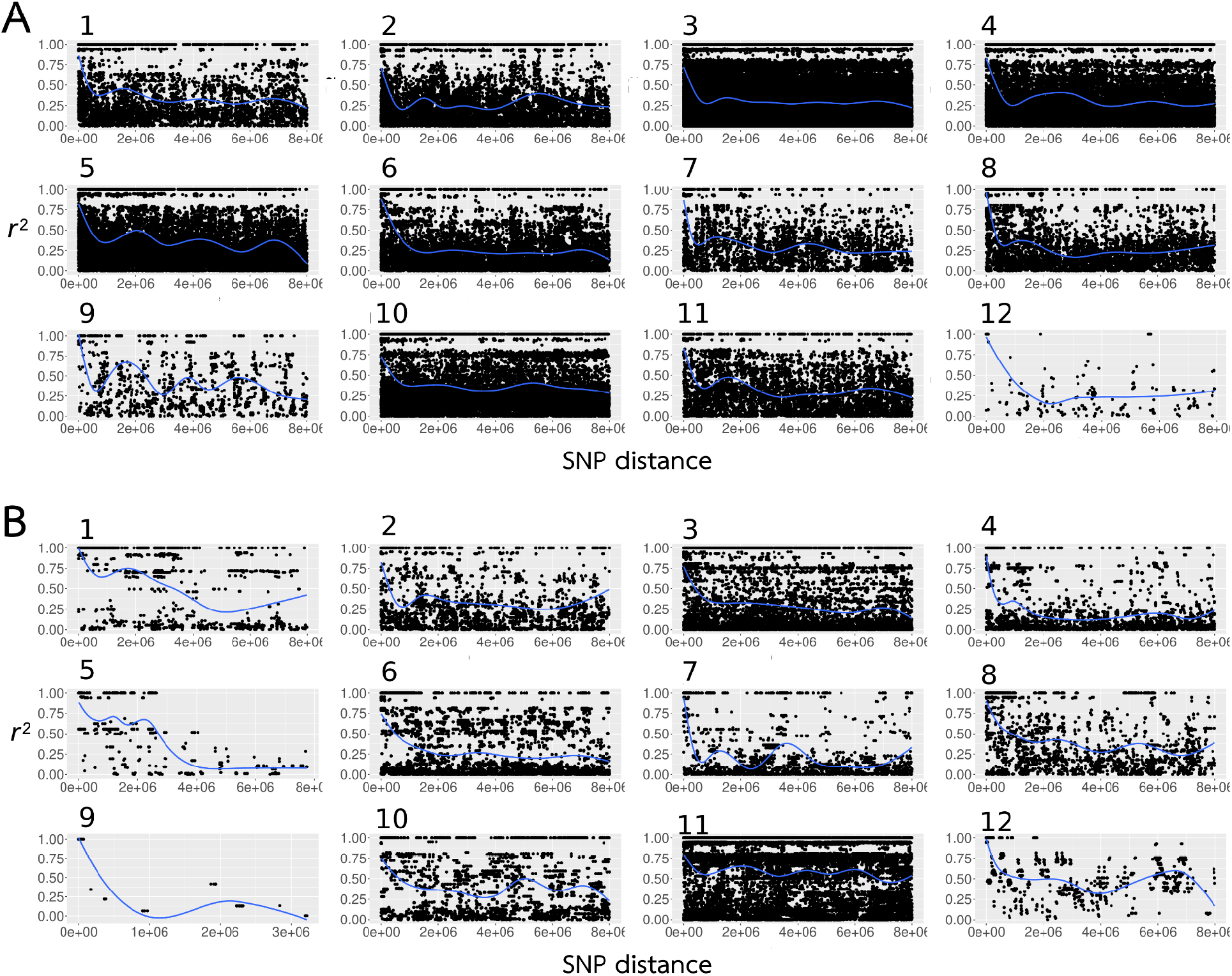
Linkage Disequilibrium (LD) decay per chromosome in parthenogenetic populations showing evidence for gene flow. LD decay was estimated for all chromosomes excluding the X, for which we did not have positional information (see methods). A line of best fit is shown in blue. **A**. *T. douglasi*, population Manchester. **B**. *T. monikensis*, population FS.

The six populations which presented no evidence for cryptic sex (hereafter obligately parthenogenetic populations), had either one clonal lineage, namely *T. genevievae –* Antonio and *T. shepardi –* Elk Mountain, or two clonal lineages, *T. genevievae –* HW20, *T. shepardi* – Spring Road, *T. douglasi* Black Bart Rd and *T. monikensis* – Sycamore (Figure 2). Even though we estimated LD values for all cases (Figures S2-9), estimates for populations with only a single clonal lineage are not informative given the extremely low polymorphism they present. Nevertheless, the presence of a single clone in those populations is in itself consistent with the expectation for the absence of sex. For the four populations with two clonal lineages, LD values within and between chromosomes were consistently very high (Figure 3 - A, Figures S2-5), indicative of a tight linkage between polymorphic positions. There was also no evidence of LD decay (Figure 3 - B, Figures S6-9), consistent with linkage between polymorphic positions and absence of recombination. Finally, individuals from all six obligately parthenogenetic populations (including the two for which LD estimates are not informative due to low polymorphism) presented consistently low heterozygosity (Figure 6), as previously shown for *Timema* individuals produced via parthenogenesis (Jaron et al. 2022; Larose et al. 2022).

**Figure 6.**
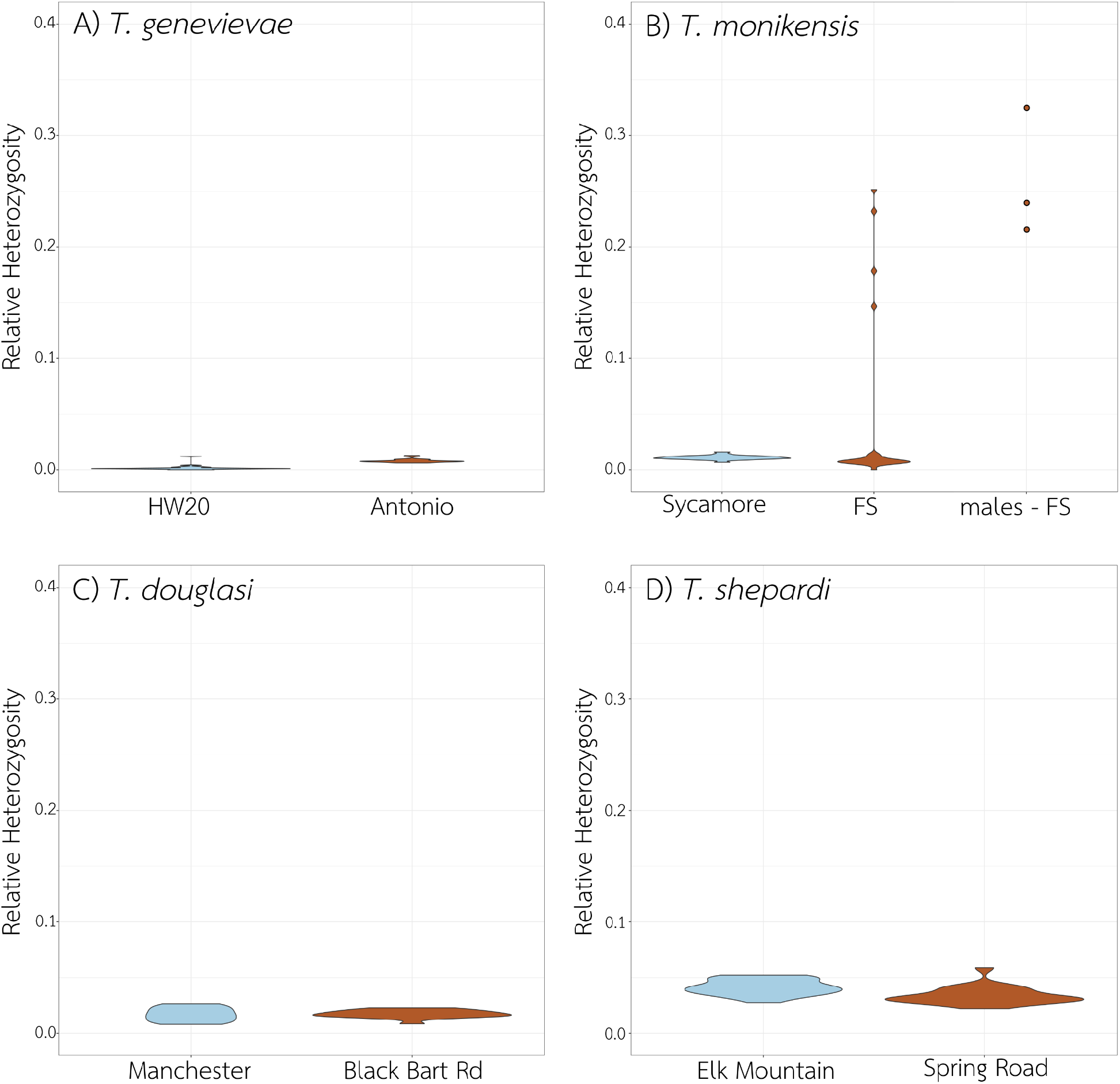
Relative heterozygosity per individual in each of the eight studied populations. **A**. *T. genevievae*, **B**. *T. monikensis*, **C**. *T. douglasi*, and **D**. *T. shepardi*. On the x axis are the populations for the four parthenogenetic species studied, plus the males in the *T. monikensis* - FS population. On the y axis is the relative heterozygosity estimated as the median between all chromosomes.

The patterns in the remaining two populations (*T. douglasi* – Manchester, and *T. monikensis –* FS) are, however, much different and best explained by (rare) sexual reproduction. Both populations present a high diversity of genotypes, with eight different clonal lineages in *T. douglasi* – Manchester, but no clonal lineages and only independent genotypes (21) in *T. monikensis* – FS (Figure 2). Regarding LD within and between chromosomes, both populations presented much lower values of *r*^2^ when compared to the strictly parthenogenetic populations (Figures 3, S2-5), yet values were still elevated relative to the sexual *T. cristinae* population (Figure 1). LD decay is also apparent in these two populations, with estimates consistently decreasing with increasing distance between markers (Figure 5). Furthermore, for *T. monikensis* – FS, *r*^2^ values within linkage groups were higher than *r*^2^ values between different chromosomes (Figure 4 - B), suggesting that segregation contributes more strongly to the high genotype diversity in this population than recombination.

Given the evidence for cryptic gene flow in the *T. douglasi* – Manchester and *T. monikensis* – FS populations, we investigated whether we could identify sexually produced individuals. Parthenogenetically produced *Timema* individuals are largely or completely homozygous (Jaron et al. 2022; Larose et al. 2022). By contrast, sex between different clones would manifest as elevated heterozygosity in the resulting offspring. In other words, the presence of individuals with elevated heterozygosity would indicate events of sex in the previous generation.

As expected, the relative heterozygosity values for the individuals in the six fully parthenogenetic populations were all very low, varying between 0 and 0.05. By contrast, four out of the 21 *T. monikensis* females from the population FS showed elevated relative heterozygosity (Figure 6), corroborating our previous conclusions of cryptic sex in this population. The heterozygosity levels of these four individuals are as expected if these individuals were produced from crosses between individuals with low heterozygosity from the same population (Figure S11). On the other hand, none of the 20 *T. douglasi* females from the Manchester population had elevated heterozygosity, indicating that none of these females were produced sexually.

Finally, to investigate whether gene flow in *T. monikensis* - FS is mediated by rare males, and to assess whether these males are produced sexually or stem from accidental X chromosome losses, we also estimated relative heterozygosity for the four available males from that population (collected in a different year). If produced sexually, males would present elevated heterozygosity, similar to the four heterozygous females, but if resultant from X loss, males would present low heterozygosity levels. All four males were heterozygotic outliers, similar to the four highly heterozygous females (Figure 6) indicating they were likely produced via sex.

To corroborate our conclusions of cryptic gene flow in the *T. douglasi* – Manchester and the *T. monikensis –* FS populations we performed three additional complementary analyses. First, we found no evidence that highly heterozygous individuals were triploid, as would be expected from the rare fertilisation of parthenogenetically produced diploid eggs (Supplemental Material, Figure S12). Second, we tested if our LD findings in the *T. monikensis -* FS population were driven by the heterozygous outliers. To do this we repeated our LD analyses with heterozygous outlier individuals removed, and found that LD and LD decay remained very similar (Figure S13). Finally, to test whether LD and heterozygosity in *T. monikensis* populations (both Sycamore and FS) were consistent between different years, we analysed additional individuals from previous collection years (namely, 2013 and 2015). We found similar results to our main analyses: no evidence for sex in Sycamore, and strong evidence for cryptic sex in the FS population from low LD (Figure S14), evident LD decay (Figure S15) and heterozygotic outlier individuals (Figure S16).

## DISCUSSION

While obligately parthenogenetic lineages are expected to be short lived, many can persist in the long term, sometimes even for millions of years (Judson and Normark 1996; Schurko et al. 2009). It is still unknown how parthenogenetic lineages manage to persist for so long and evade the costs of lacking sex. However, rare events of sexual reproduction have been suggested as an explanation for the persistence of long lived parthenogens, and evidence for rare gene flow in “obligately” parthenogenetic species is accumulating (Signorovitch et al. 2015; Vakhrusheva et al. 2020; Boyer et al. 2021; Laine et al. 2022).

Rare events of sex in obligately parthenogenetic species can be difficult to demonstrate, since unless we find direct evidence (such as direct observations of sexual offspring in broods of parthenogenetic females), we need to rely on indirect evidence. Indirect evidence may include the presence of functional males, unexpectedly high genetic diversity in parthenogenetic populations or signatures of recombination (e.g. recombining haplotypes, linkage disequilibrium). Here, we searched for signatures of recombination and sex in parthenogenetic species of the genus *Timema* by estimating genotype diversity within populations, linkage disequilibrium (LD), LD decay with increasing marker distance, and heterozygosity. Some forms of parthenogenesis can involve recombination and segregation (White 1973; Engelstädter 2017), but in homozygous parthenogens such as *Timema*, these two mechanisms have little to no effect on genotype diversity. In such homozygous parthenogens, signatures of recombination can therefore provide very strong evidence for sexual reproduction.

While we found no evidence for cryptic sex in six populations, we detected signatures of recombination and cryptic sex in two populations of parthenogenetic *Timema*, the population Manchester of *T. douglasi* and the population FS of *T. monikensis*. Cryptic sex, at least in *T. monikensis*, is almost certainly mediated by rare males, known to occur in *Timema* parthenogens. However, the uncovered patterns in the Manchester and FS populations reflected different scenarios. Regarding the Manchester population (*T. douglasi*), LD within and between chromosomes was generally low (and similar), and a clear pattern of LD decay was evident across all linkage groups. Genotype diversity was relatively high (i.e., we found eight different “clones” among the 20 genotyped individuals), and clones were often represented by multiple individuals (1-5 per clone). Finally, the heterozygosity of all females was as low as in the strictly parthenogenetic populations, indicating they were produced via parthenogenesis. These findings are best explained by rare sexual events that occurred in the past, leading to a high diversity of genotypes, but also allowing for the spread of these genotypes into clonal lineages.

The second population with evidence for gene flow, the FS population of *T. monikensis*, likely features more recent sexual events than the Manchester population of *T. douglasi*. The 21 genotyped females represented 21 distinct genotypes, as would be expected for a sexual but not for a largely parthenogenetic population. Consistent with the high genotype diversity, LD decay was evident for all chromosomes and LD values were generally very low. Finally, 4 out of the 21 females (∼19%) were characterised by elevated heterozygosity, and were most likely produced from a sexual cross between different genotypes (Figure S6). *Timema* parthenogenetic females were previously screened for their ability to fertilise eggs by using crosses with males from related sexual species (*T. cristinae* males for *T. monikensis* females of the FS population), but none of the 322 genotyped hatchlings were sexually produced (Schwander et al. 2013). Even though the absence of sexually produced offspring in those crosses could partly stem from species divergence and associated incompatibilities, the reciprocal crosses (*T. cristinae* females mated to *T. monikensis* males) result in large numbers of hybrid offspring (Schwander et al. 2013). Given our new results indicating *T. monikensis* are capable of reproducing sexually, this suggests that sexually produced offspring were too rare to be detected in the analysed sample of hatchlings, yet we here report that as many as 19% of the adult females of the FS population were sexually produced. One possible explanation for such a frequency difference could be that sexually produced females have a survival advantage over parthenogenetically produced ones. If that is the case, then the maintenance of mostly parthenogenetic reproduction in the FS population in spite of this apparent selection for sex will be an interesting focus for future research.

Even though we do find evidence for genetic exchange in two parthenogenetic populations, most of the populations analysed here are *de facto* parthenogenetic, and do not present any evidence of gene flow (present or recent past). Two scenarios could potentially explain this patchy occurrence of sex. In the first scenario, males appear spontaneously in a parthenogenetic all-female population (via e.g. X aneuploidies), and parthenogenetic females were then able to mate and produce at least some sexual offspring. Such sexual offspring would comprise males, which would then allow for the propagation of males in the population. In the second scenario, males and sexual reproduction were originally present in all populations, but lost in the majority of them. Why sex occurs in some populations but not others is unclear and may also be a dynamic process. For example, if sex can emerge spontaneously in a population it may persist for multiple generations before going extinct due to the short-term advantages of parthenogenesis. In species that inhabit fire prone habitats, such as *Timema*, an ecologically important short-term advantage of parthenogenesis is the ability to rapidly recolonise areas following fire with a single (female) individual. As such, the recurrent selection for parthenogenesis following fire, combined with the inherent stochasticity of extinction-recolonisation dynamics of fire-prone habitats, may drive a patchy distribution of sex in *Timema* parthenogenetic populations.

As mentioned above, males are known in four out of the five described parthenogenetic species of *Timema*, including *T. monikensis* (Sandoval and Vickery 1996; Vickery and Sandoval 1999, 2001; Schwander et al. 2013). Our findings of cryptic sex in one population of *T. monikensis* now raise the question of whether males in other *Timema* species could also mediate cryptic sex in some populations, albeit much more rarely than in the FS population of *T. monikensis*. In combination with other recent reports of gene flow in parthenogenetic animal species (Signorovitch et al. 2015; Vakhrusheva et al. 2020; Boyer et al. 2021; Laine et al. 2022) our findings suggest that episodes of rare sex in largely parthenogenetic species may occur more frequently than generally appreciated. Thus, it would be useful to screen additional parthenogenetic species for signatures of sex, especially species with only suggestive evidence that sexual reproduction may still be ongoing, such as parthenogenetic species with functional males, or unexpectedly high levels of genetic diversity. Males have been found in several parthenogenetic species (e.g. (Simon et al. 1991; Butlin et al. 1998; Delmotte et al. 2001; Tvedte et al. 2020)), and many examples exist for high genetic diversity in parthenogenetically reproducing populations. Both these findings have been used to suggest cryptic gene flow in parthenogenetic species (Belshaw et al. 1999; Ross et al. 2013; Fontcuberta et al. 2016), but neither are reliable evidence for sex. The presence of males itself may not necessarily mean that those males can mate with parthenogenetic females, since traits required for sexual reproduction are typically vestigialised in parthenogenetic females (van der Kooi and Schwander 2014). Furthermore, males can be dysfunctional, especially in older parthenogenetic lineages where mutations may have accumulated in pathways for exclusively male functions (van der Kooi and Schwander 2014). Finally, males produced by parthenogenetic females could mate with sexual females, such as in the cases of contagious parthenogenesis (Simon et al. 1999; Delmotte et al. 2001). Accordingly, high genetic diversity in parthenogenetic species could be explained by multiple transitions to parthenogenesis (Fontaneto et al. 2008; Janko et al. 2012; Capel et al. 2017) and not indicate cryptic sex. However, future studies of such species could reveal cryptic sex in at least some of them. Finally, despite the evidence for cryptic sex in some parthenogenetic species, evidence for long term parthenogenesis in eukaryotes has also been reported for protozoans (Weir et al. 2016) and oribatid mites (Brandt et al. 2021). Since both these groups are characterised by extremely large population sizes, these findings could be an indication of support for theory showing diminishing benefits of sex with large population size (Gabriel et al. 1993; Gordo and Charlesworth 2000).

In conclusion, independently of how many additional parthenogenetic species will be discovered to have rare sex, the fact that at least some presumed obligate parthenogens can reproduce sexually questions our understanding about the plasticity of sexual and parthenogenetic reproduction modes in animals. When classifying a species as sexual, we expect that all offspring is produced by sex, yet many species are capable of spontaneous parthenogenesis (White 1973). Accordingly, when classifying a species as obligately parthenogenetic we rarely consider occasional and/or non-canonical sex. While most reproductive systems in animals fit into a bimodal pattern of “sexual” or “parthenogenetic”, we also concur with (Boyer et al. 2021), that reproductive systems may contain more variation than previously assumed, and believe this could in many ways explain how old parthenogens escaped the long term costs of the “lack of sex”.

## Acknowledgements

The authors thank Roger K. Butlin for his constructive comments to the methods used here, and Chloe Larose and Bart Zjilstra for support during field work. Most analyses were run using the computing infrastructure at the University of Lausanne maintained by the former Vital-IT platform of the SIB Swiss Institute of Bioinformatics (SIB) and the DCSR. We would like to acknowledge funding from the European Research Council (Consolidator Grant No Sex No Conflict) and Swiss FNS grant 31003A_182495 to TS.

## Data and code availability

Raw sequence reads have been deposited in NCBI’s sequence read archive under the bioproject: PRJNA798556. Scripts for the analyses in this paper are available at: https://github.com/SusanaNFreitas/cryptic_gene_flow.

