## Supplemental Material for "Evidence for cryptic gene flow in parthenogenetic stick insects of the genus *Timema*"

#### Corroborating cryptic sex in two parthenogenetic populations

In order to corroborate our inference of cryptic sex from patterns of LD and the heterozygotic outliers in two populations (*T. douglasi* – Manchester, and *T. monikensis* – FS), we conducted two additional analyses. First, we tested if heterozygous individuals could be triploid, which may be the case if diploid parthenogenetic eggs get fertilised by a haploid sperm. In a diploid individual both alleles at a heterozygous site are expected to have equal coverage, whereas in a triploid individual one allele is expected to have twice the coverage. Estimated minor:major allele depth ratio distributions, expected to be skewed to 1/2 for a triploid individual, and skewed to 1 for a diploid individual, did not suggest that heterozygous individuals are triploid (Figure S12).

Second, we repeated all LD estimations in the FS population after excluding the heterozygous outlier individuals, and the results obtained were indistinguishable to the same analyses with all samples. This means that the heterozygous individuals are not driving our results (Figure S13).

### Supplemental Figures

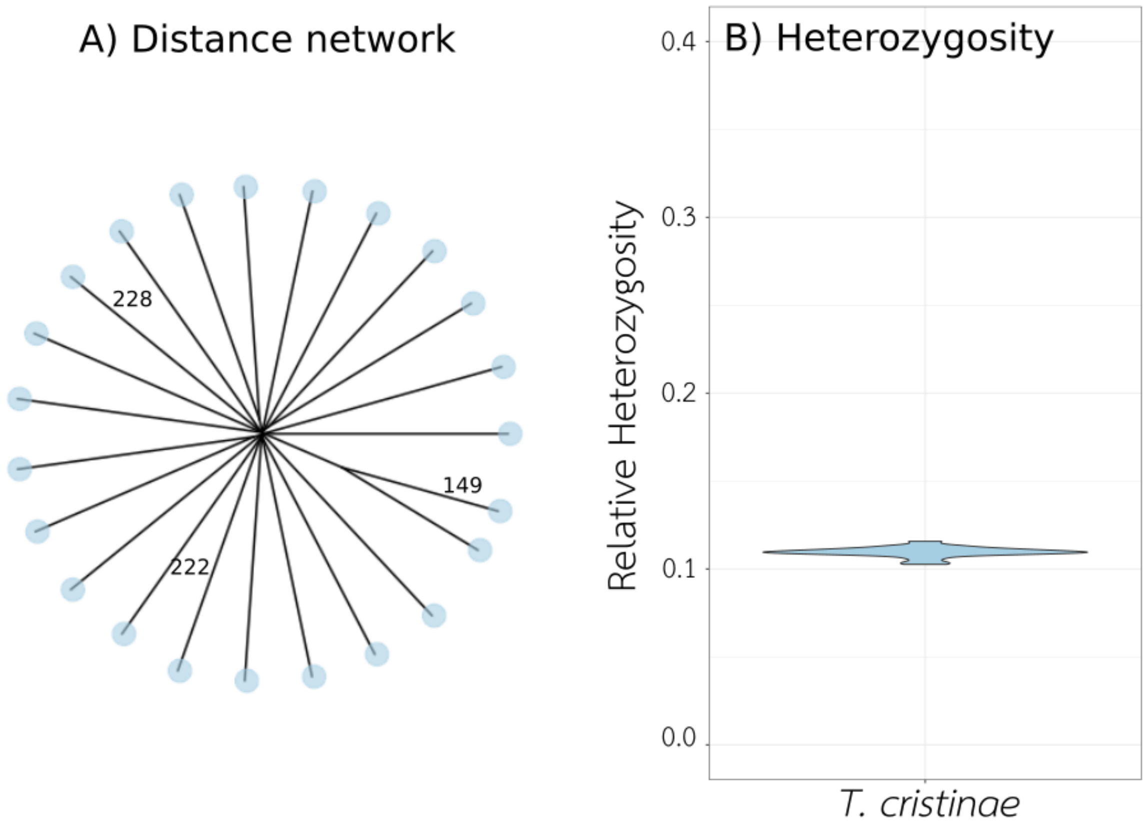


Figure S1 - High genotype diversity (A) illustrated by pairwise genetic distances between individuals (numbers on branches indicate Euclidean distance), and relative heterozygosity (B) (proportion of heterozygous SNPs among polymophic sites) for the sexual species *T. cristinae*.

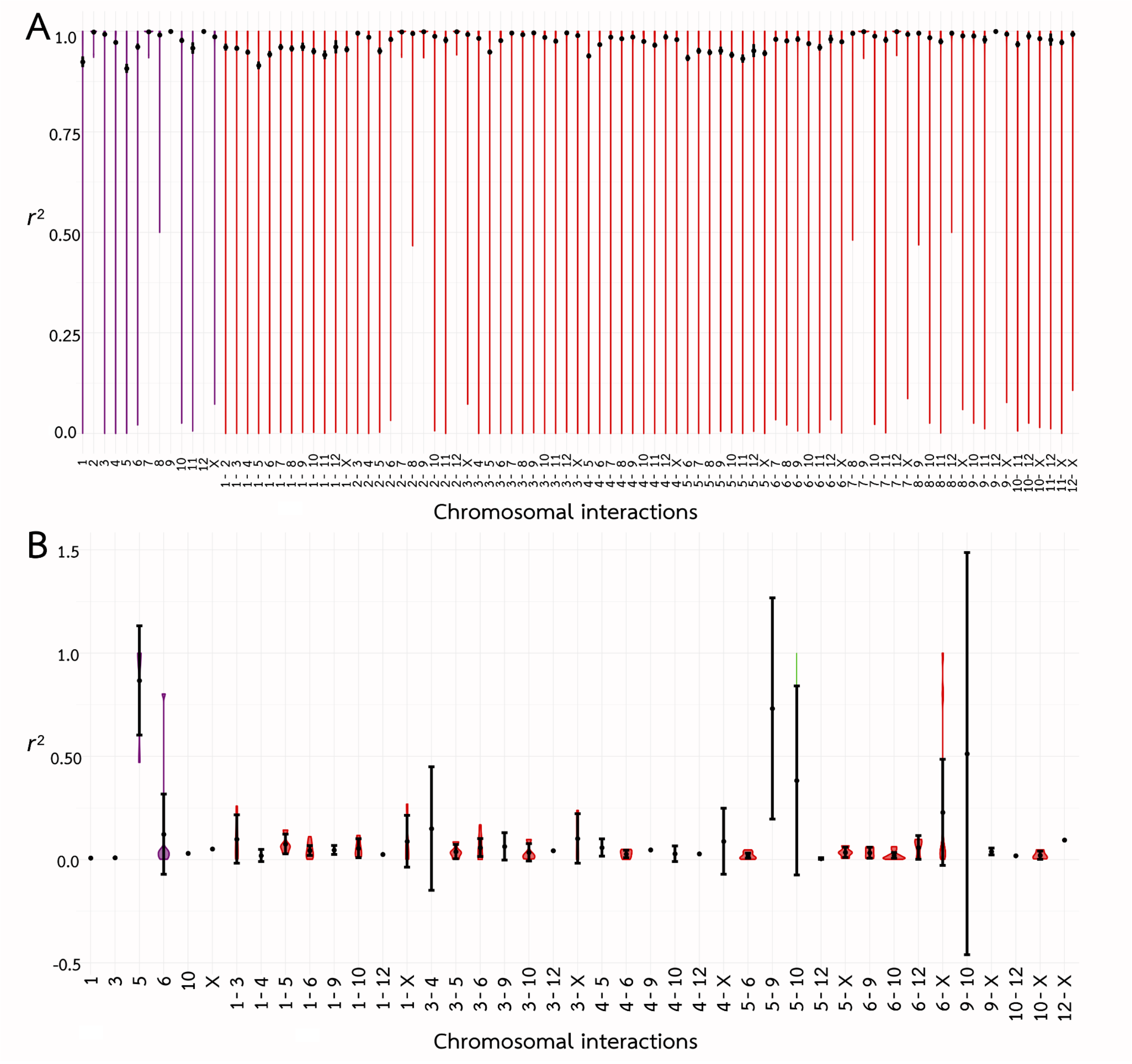


Figure S2 - Linkage Disequilibrium (LD) within and between chromosomes (linkage groups) estimated with $r^2$ for the parthenogenetic species *T. genevievae*. In panel A, population HW20, and in panel B population Antonio (where LD estimates are unreliable due to too little polymorphism among individuals of a single clone, see Figure 2-A). In both panels the x axis corresponds to the chromosome interactions, in purple SNPs within the same chromosome, and in red SNPs between different chromosomes, and the y axis to the LD ($r^2$) values.

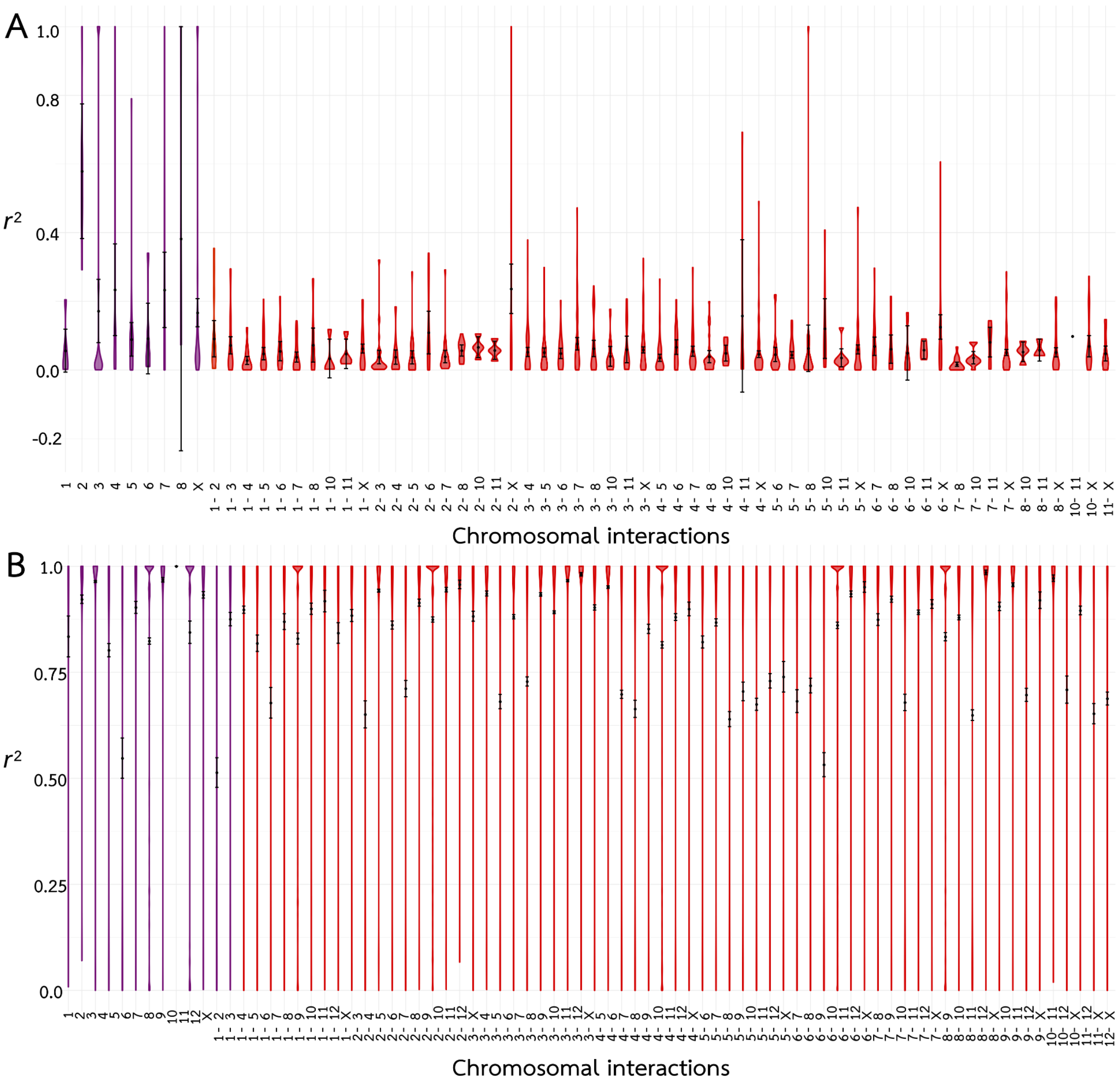


Figure S3 - Linkage Disequilibrium (LD) within and between chromosomes (linkage groups) estimated with $r^2$ for the parthenogenetic species *T. shepardi*. In panel A, population Elk Mountain (where LD estimates are unreliable due to too little polymorphism among individuals of a single clone, see Figure 2-D; only chromosomes with SNPs are plotted), and in panel B population Spring Road. In both panels the x axis corresponds to the chromosome interactions, in purple SNPs within the same chromosome, and in red SNPs between different chromosomes, and the y axis to the LD ($r^2$) values.

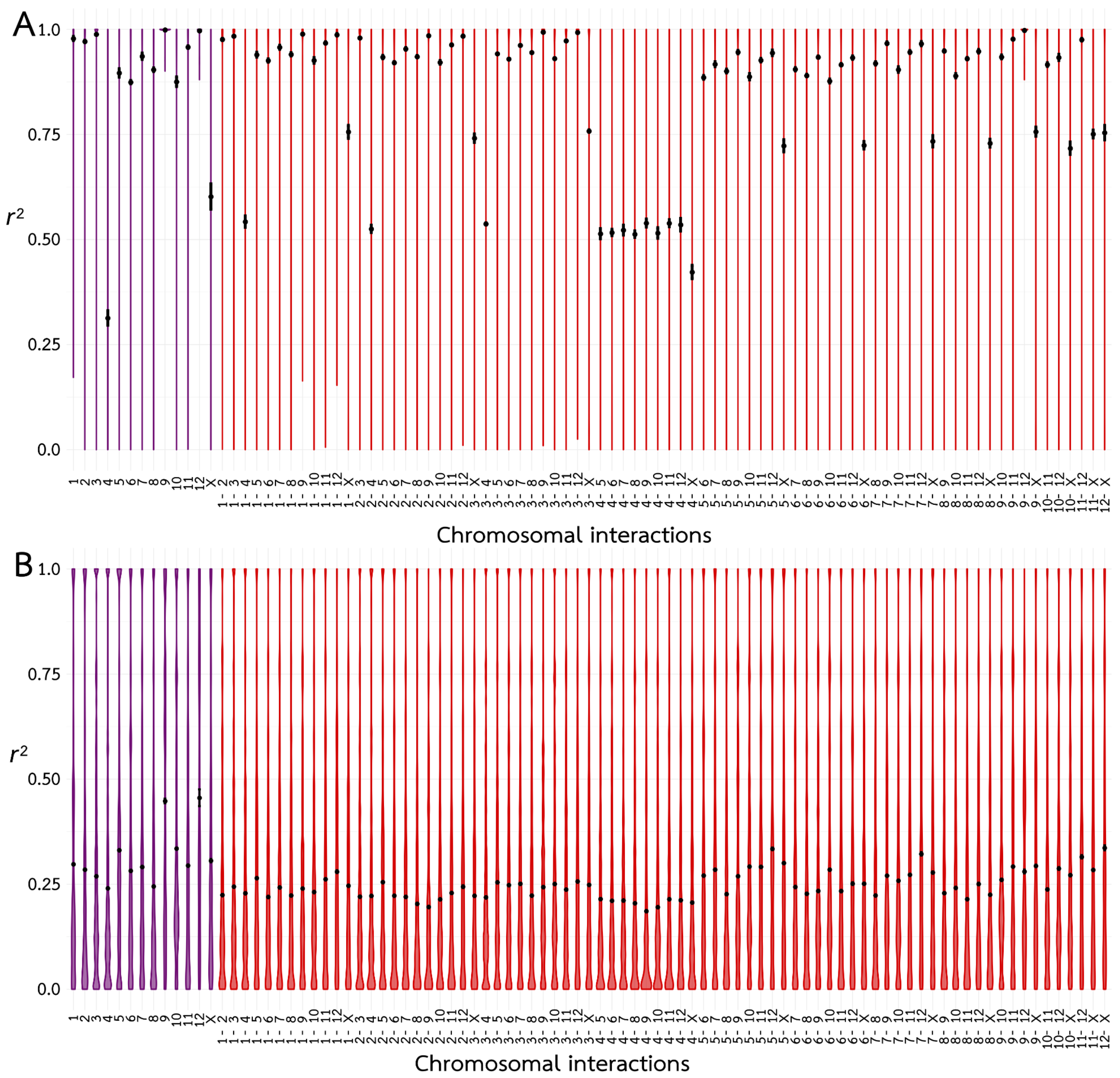


Figure S4 - Linkage Disequilibrium (LD) within and between chromosomes (linkage groups) estimated with $r^2$ for the parthenogenetic species *T. douglasi* (panel A - population Black Bart Road, panel B population Manchester). In both panels the x axis corresponds to the chromosome interactions, in purple SNPs within the same chromosome, and in red SNPs between different chromosomes, and the y axis to the LD ($r^2$) values (calculated for all chromosomes).

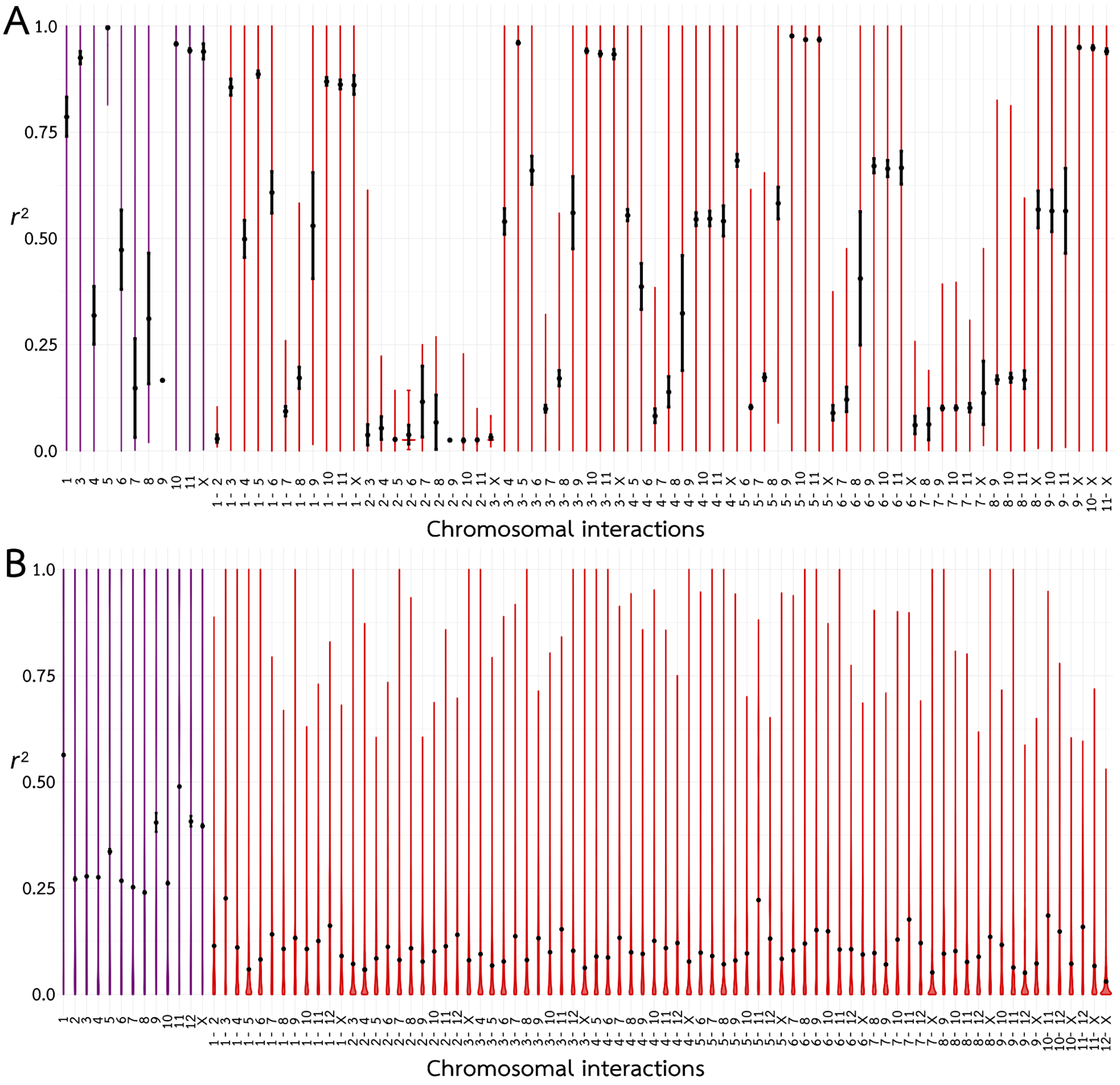


Figure S5 - Linkage Disequilibrium (LD) within and between chromosomes (linkage groups) estimated with $r^2$ for the parthenogenetic species *T. monikensis* (panel A - population Sycamore, panel B - population FS). In both panels the x axis corresponds to the chromosome interactions, in purple SNPs within the same chromosome, and in red SNPs between different chromosomes, and the y axis to the LD ($r^2$) values (calculated for all chromosomes).

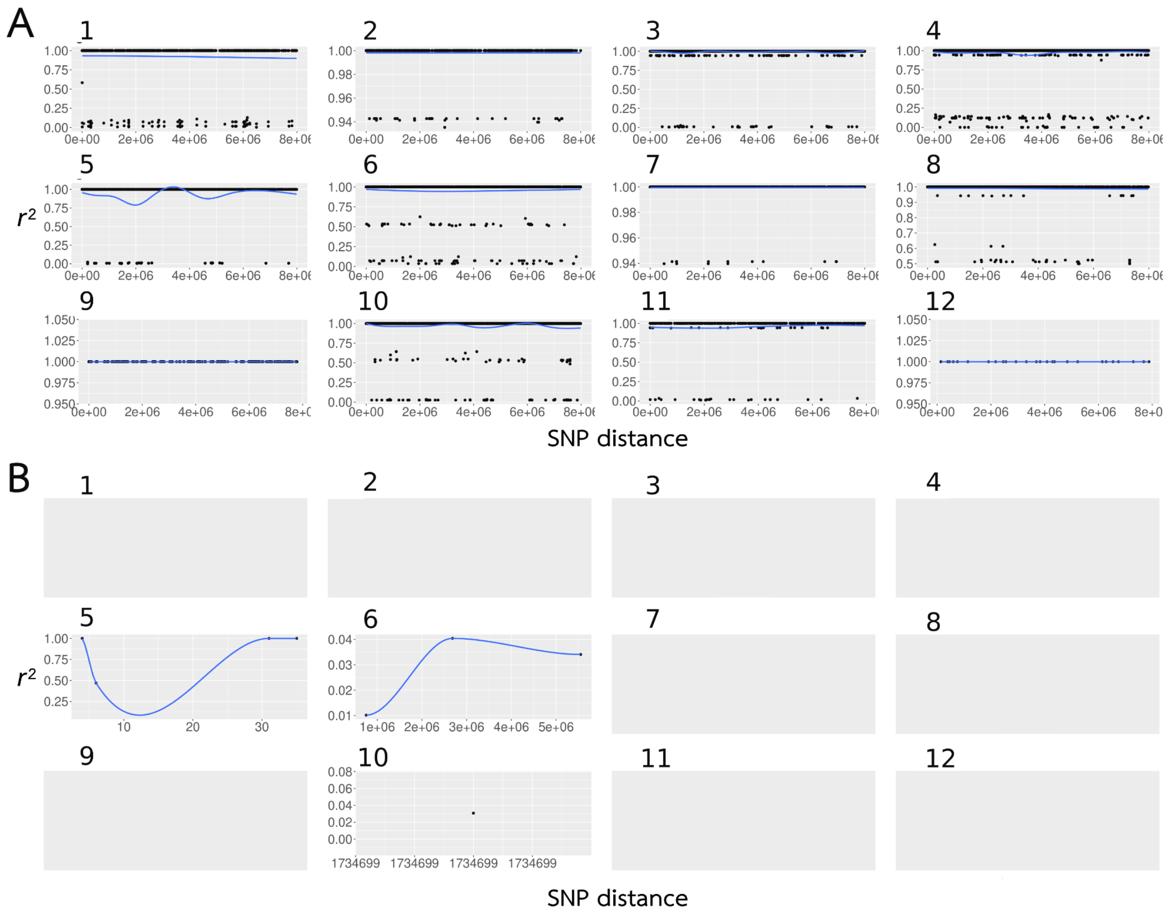


Figure S6 - LD decay per chromosome (linkage group 1-12) estimated with $r^2$ for the parthenogenetic species *T. genevievae.* LD decay was calculated for all chromosomes excluding the X (for which we did not have positional information, see methods). On the y axis are the $r^2$ values, on the x axis the distance between SNPs. Panel A corresponds to population HW20, panel B - population António (data are plotted only for chromosomes with available SNPs).

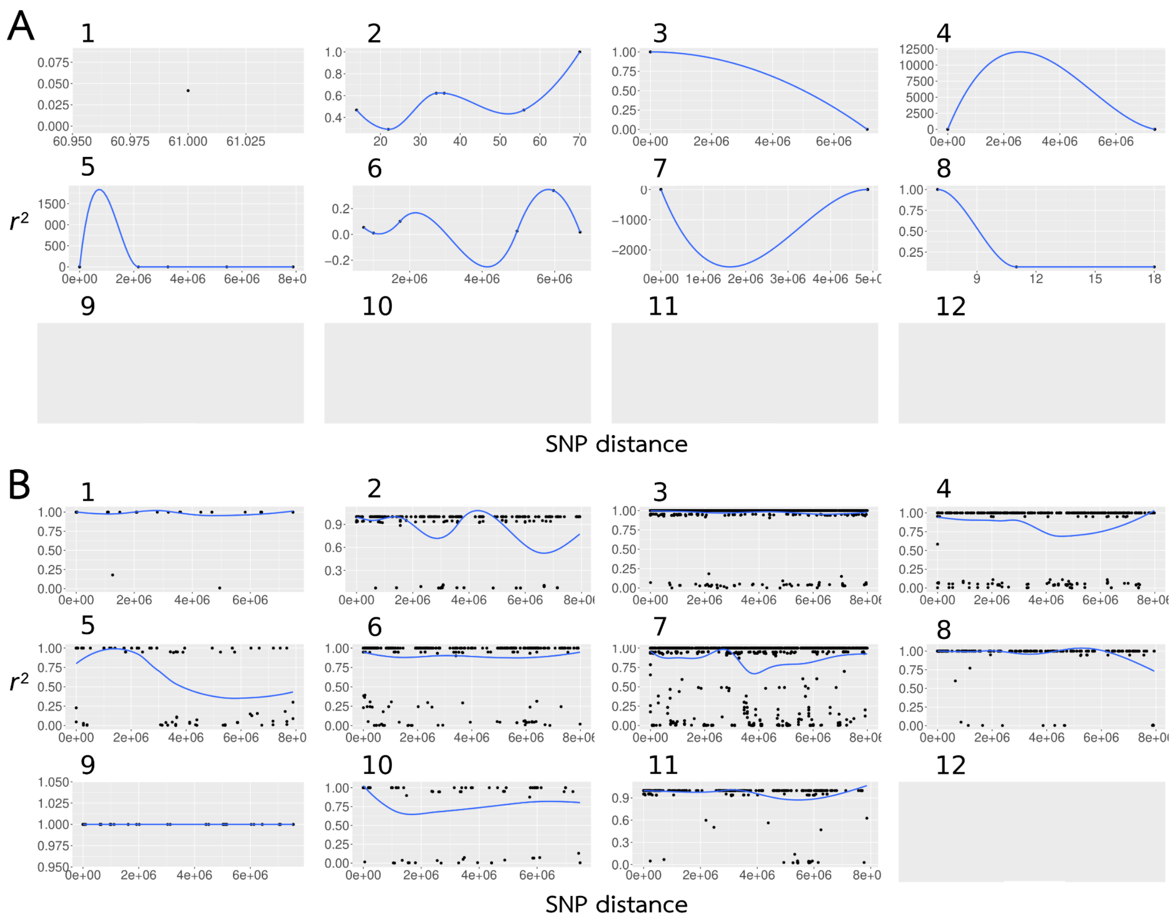


Figure S7 - LD decay per chromosome (linkage group 1-12) estimated with $r^2$ for the parthenogenetic species *T. shepardi.* On the y axis are the $r^2$ values, on the x axis the distance between SNPs. LD decay was calculated for all chromosomes excluding the X (for which we did not have positional information, see methods). Panel A - population Elk Mountain, panel B - population Spring Road (data are plotted only for chromosomes with available SNPs).

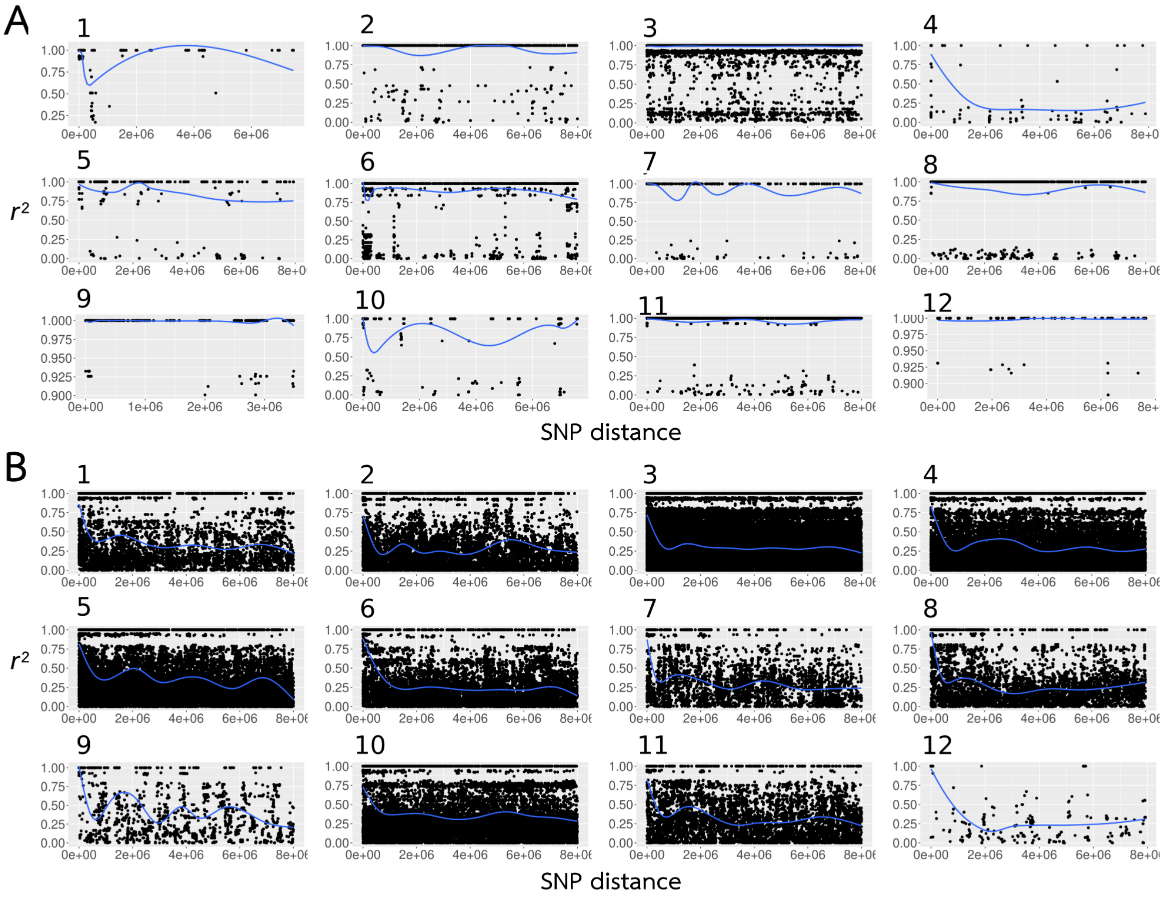


Figure S8 - LD decay per chromosome (linkage group 1-12) estimated with $r^2$ for the parthenogenetic species *T. douglasi.* On the y axis are the $r^2$ values, on the x axis the distance between SNPs. LD decay was calculated for all chromosomes excluding the X (for which we did not have positional information, see methods). Panel A - population Black Bart Rd, panel B - population Manchester.

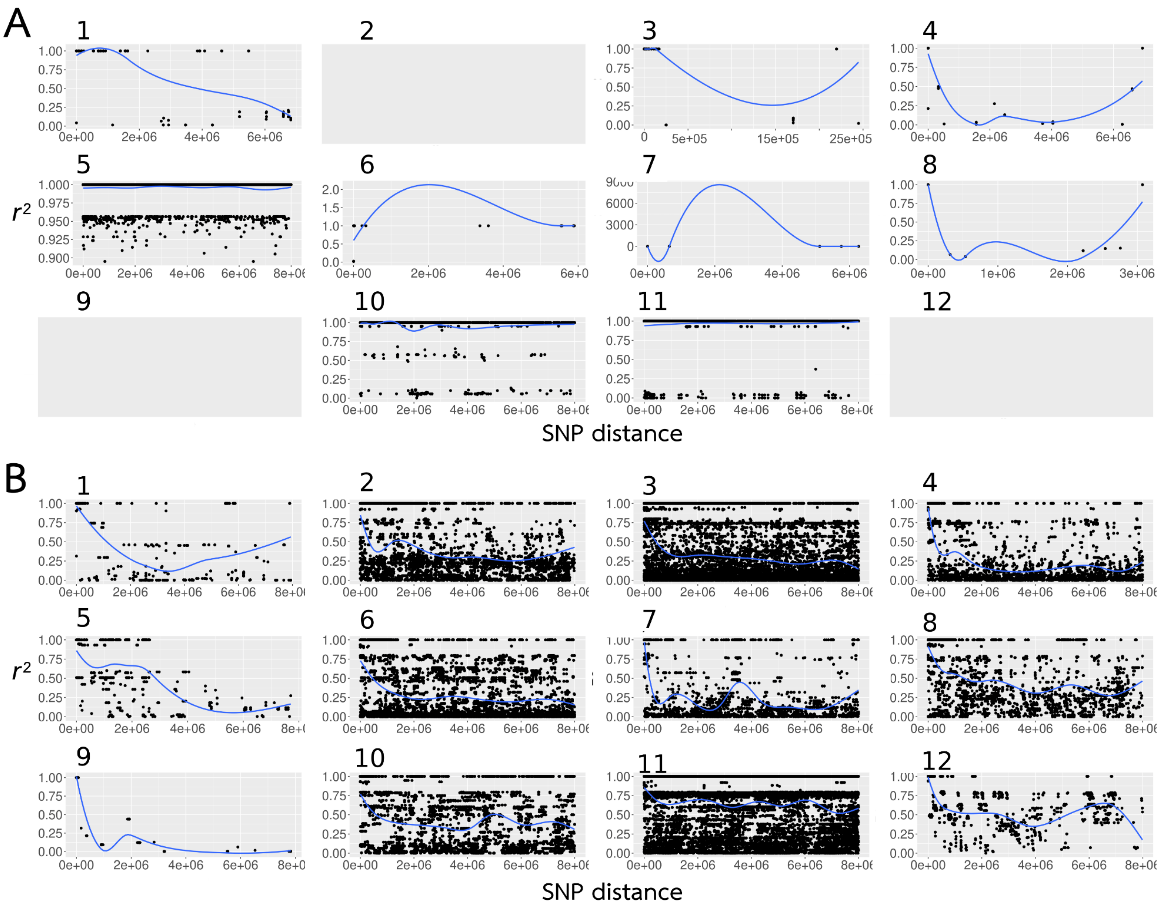


Figure S9 - LD decay per chromosome (linkage group 1-12) estimated with $r^2$ for the parthenogenetic species *T. monikensis.* On the y axis are the $r^2$ values, on the x axis the distance between SNPs. LD decay was calculated for all chromosomes excluding the X (for which we did not have positional information, see methods). Panel A - population Sycamore, panel B - population FS (data are plotted only for chromosomes with available SNPs).

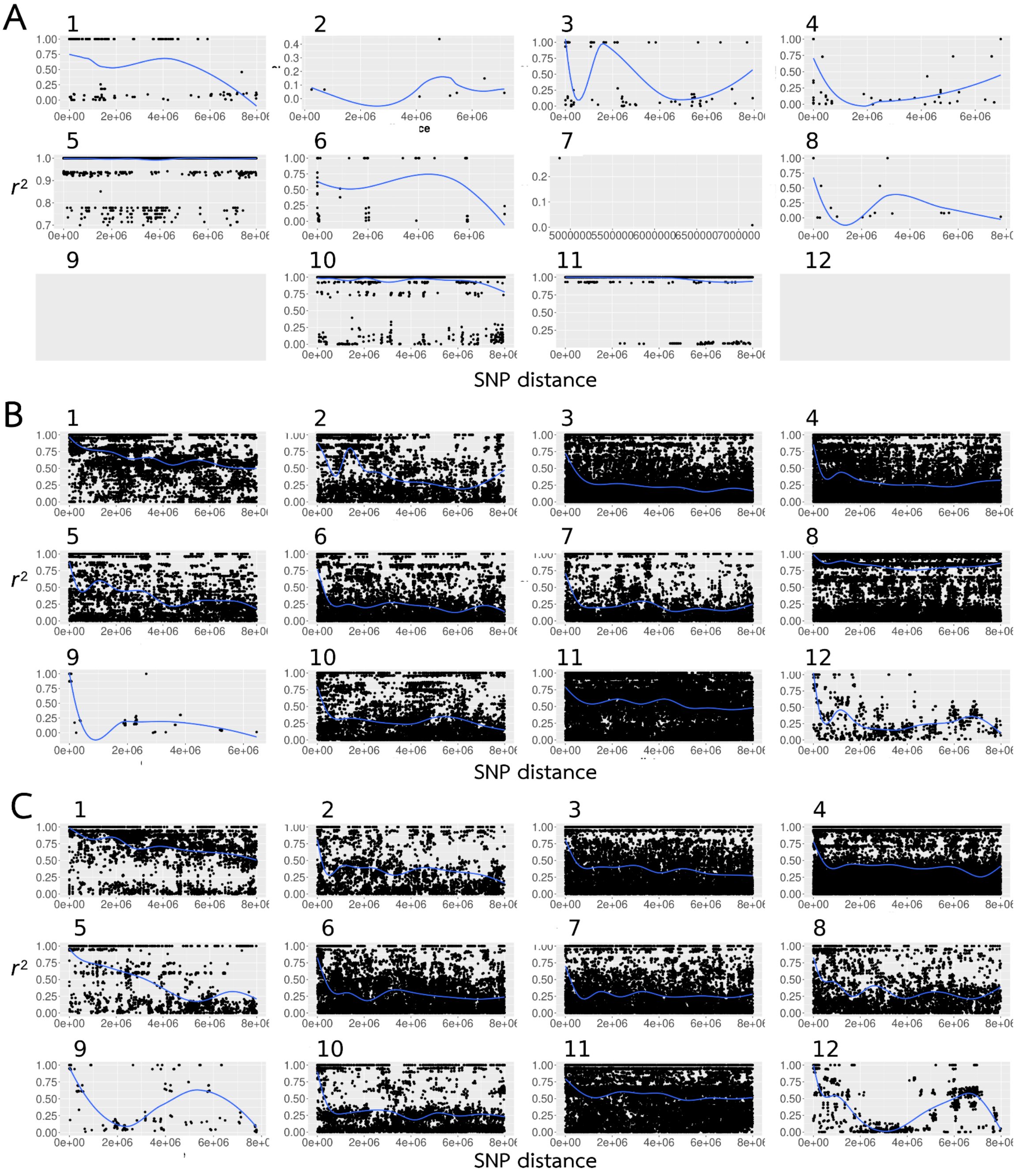


Figure S10 - LD decay per chromosome (linkage group 1-12) estimated with $r^2$ for the additional individuals from the parthenogenetic species *T. monikensis.* On the y axis are the $r^2$ values, on the x axis the distance between SNPs. LD decay was calculated for all chromosomes excluding the X (for which we did not have positional information, see methods). Panel A - population Sycamore (sampling year 2013), panel B -population FS (2013), and panel C- population FS (2015) (data are plotted only for chromosomes with available SNPs).

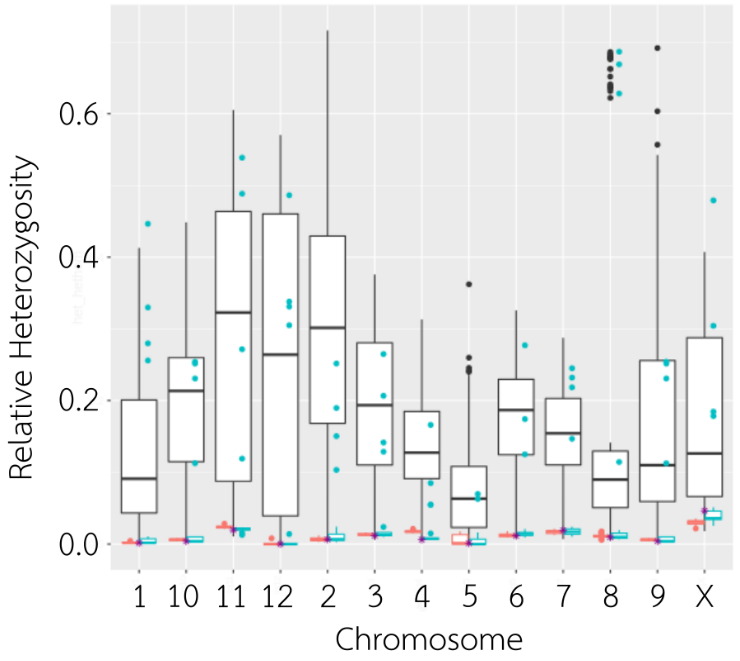


Figure S11 - Relative heterozygosity of *T. monikensis* females from FS (in blue) and Sycamore (in coral, added for reference) populations, depicted separately for each chromosome. White bars correspond to the expected heterozygosity of putative crosses between individuals (non-outliers) from the FS population. Expected heterozygosity was estimated using only homozygous positions in all available random pairs of individuals. When one SNP was homozygous for a different allele between the two individuals, it was counted as a heterozygous position in the offspring, and accordingly when that SNP was homozygous for the same allele in the different individuals, it was counted as an homozygous in the offspring. Relative heterozygosity was then estimated as the ratio between heterozygous positions in the offspring over the total number of SNPs used.

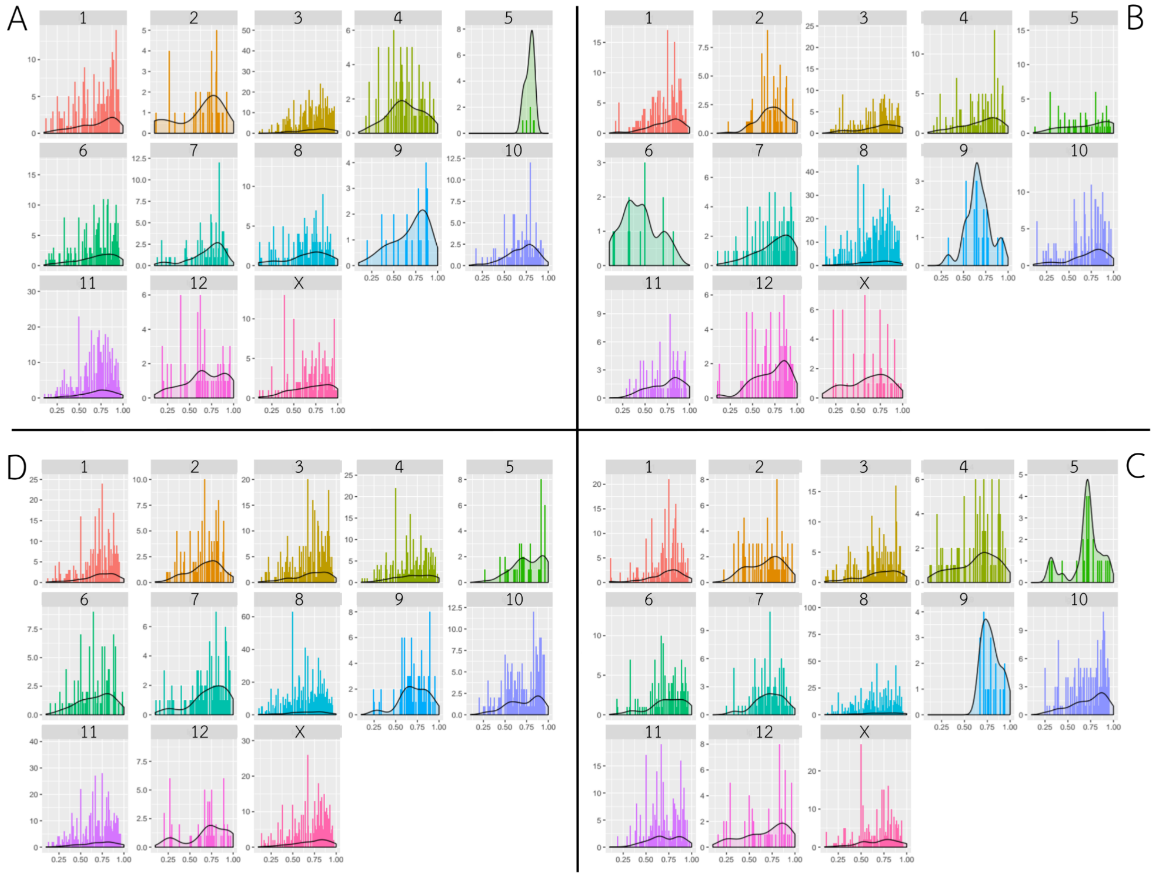


Figure S12 - Distribution of minor allele by major allele read depth ratio per chromosome (linkage groups 1-12, and the X chromosome) for the four highly heterozygous *T. monikensis* females from FS: samples A) Tms1_P2, B) Tms17_P2, C) Tms22_P2, and D) Tms19_P2 (see Table S1). The fitted line in each histogram corresponds to the smoothed version of the distribution (kernel density estimate obtained with the geom_density() function available from the ggplot2 R package). In diploid individuals the distribution is expected to be skewed to ~1 (but <1 because of sampling variance), whereas in triploid individuals it is ~0.5.

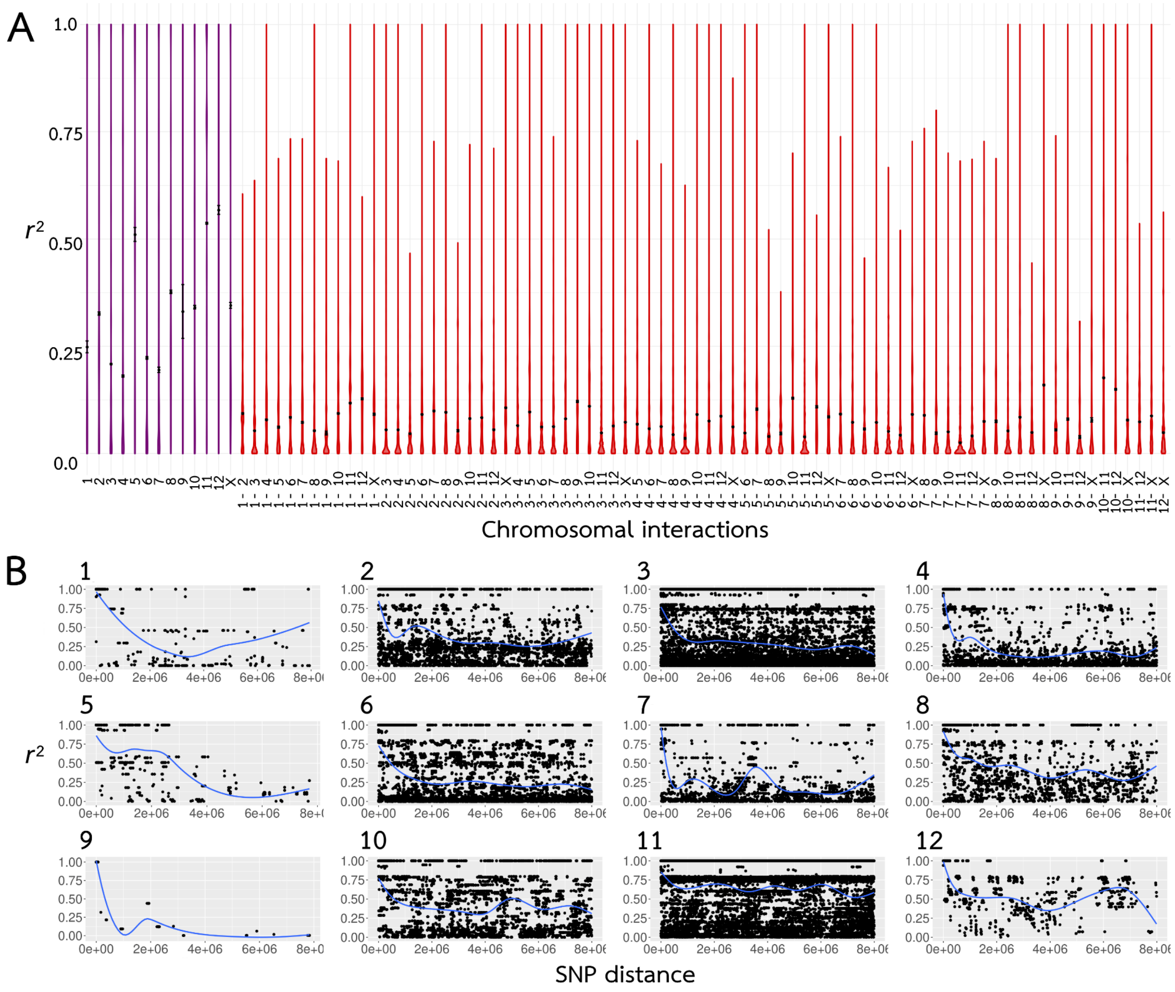


Figure S13 - A) $r^2$ (LD) values within (purple) and between (red) chromosomes and B) LD decay for the *T. monikensis* - FS population with cryptic gene flow, excluding the four heterozygous outlier females. On the y axis are the $r^2$ values, on the x axis the distance between SNPs.

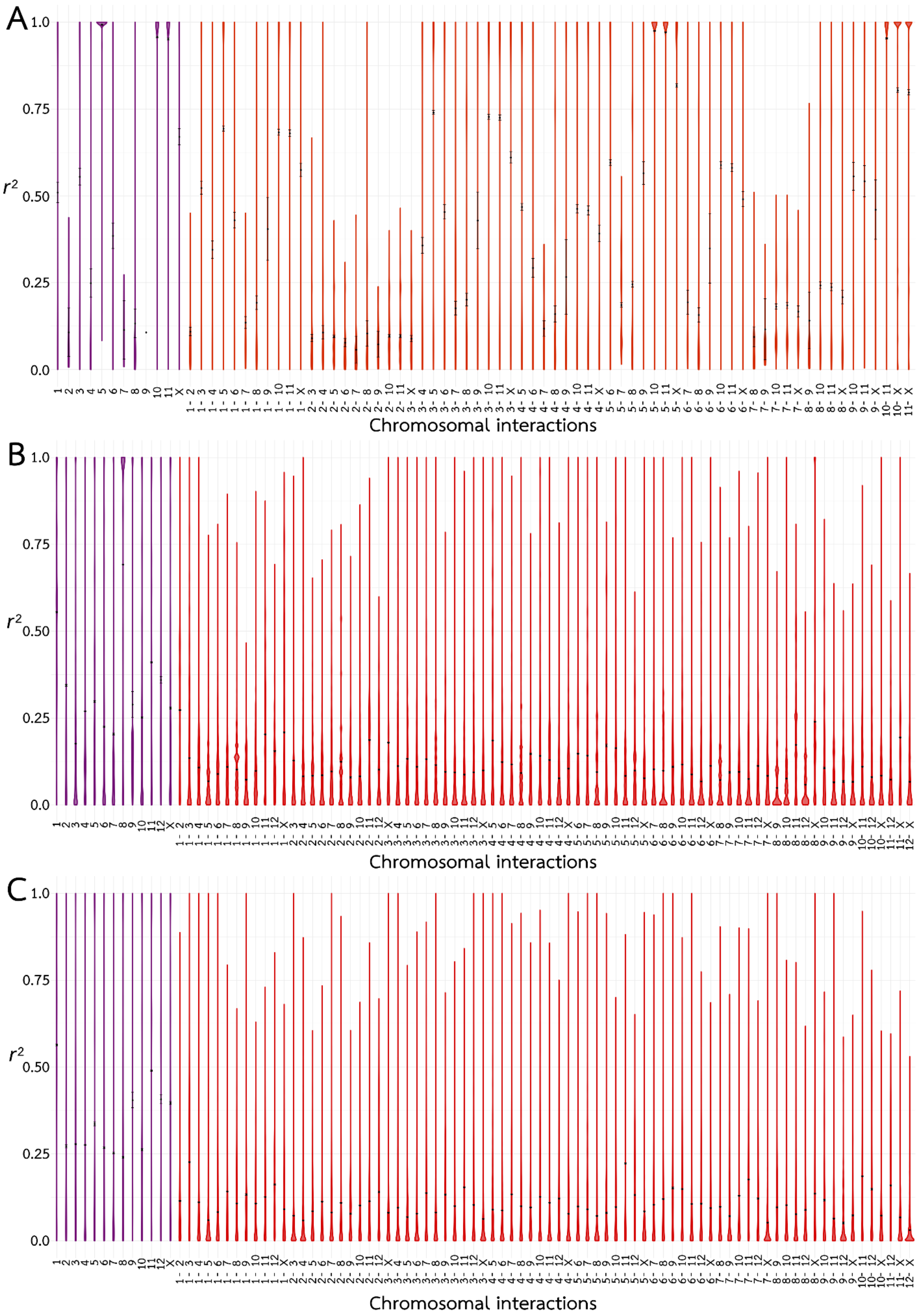

A
1.0
0.75
r2 0.50
0.25
0.0
Chromosomal interactions
B
1.0
0.75
r2 0.50
0.25
0.0
Chromosomal interactions
C
1.0
0.75
r2 0.50
0.25
0.0
Chromosomal interactions

Figure S14 - $r^2$ (LD) values within (purple) and between (red) chromosomes for the additional *T. monikensis* individuals (not included in the first analyses) collected from A) Sycamore (sampling year 2013), and B) FS (2013) and C) FS (2015).

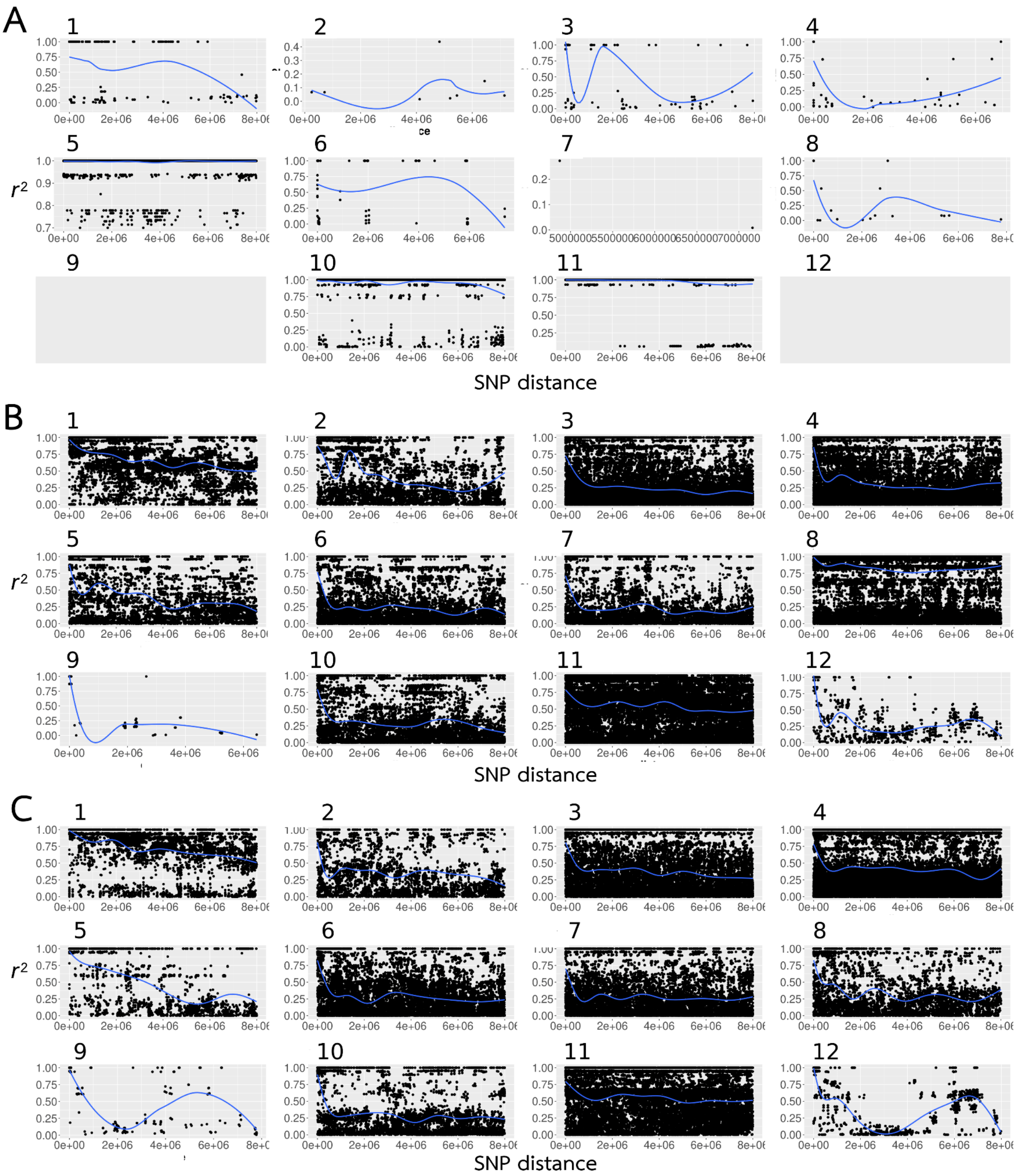

Figure S15 - LD decay values ($r^2$) within increasing distances in the chromosomes for the additional *T. monikensis* individuals (not included in the first analyses) collected from A) Sycamore (sampling year 2013), and B) FS (2013) and C) FS (2015). On the y axis are the $r^2$ values, on the x axis the distance between SNPs. Data are plotted only for chromosomes with available SNPs.

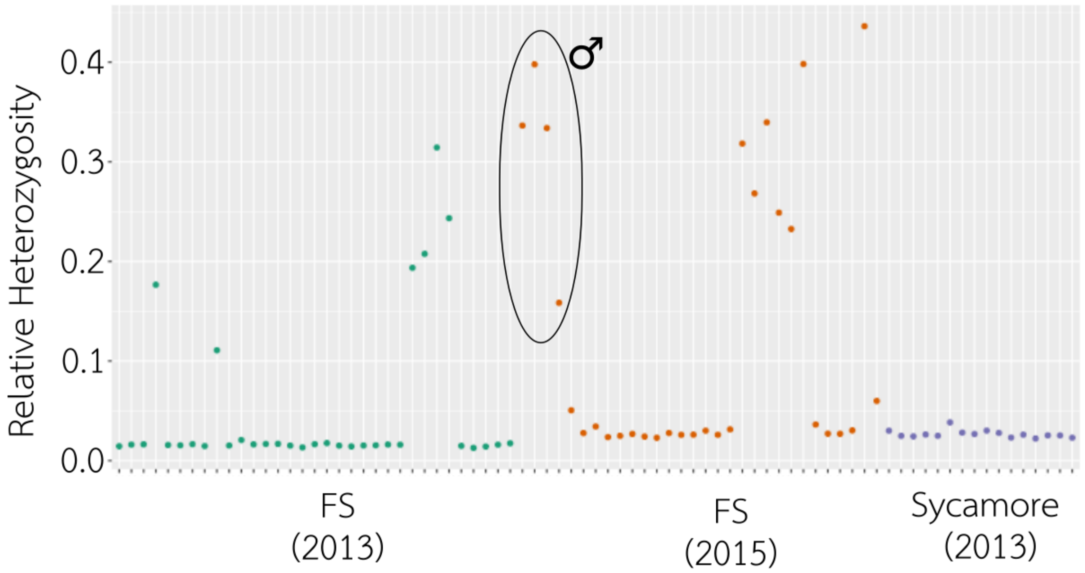


Figure S16 - Relative heterozygosity of the additional *T. monikensis* individuals used to corroborate cryptic gene flow in the FS population (green and orange distinguish individuals collected in 2013 and 2015, respectively), and the absence thereof in the Sycamore population (in violet, collected in 2013). The frequency of sexually produced females in the FS population is similar in both sampling years (6 out of 33 in 2013; 7 out of 26 in 2015; p=0.53). Male *T. monikensis* were included here (in orange, from the 2015 collection), but were not included in the frequency of sexually produced offspring, since they were not randomly sampled. For detailed sample information see Table S1.
